# Genetic comorbidity between major depression and cardio-metabolic disease, stratified by age at onset of major depression

**DOI:** 10.1101/645077

**Authors:** SP Hagenaars, JRI Coleman, S Choi, H Gaspar, MJ Adams, D Howard, K Hodgson, M Traylor, TM Air, TFM Andlauer, V Arolt, BT Baune, EB Binder, DHR Blackwood, DI Boomsma, A Campbell, M Cearns, D Czamara, U Dannlowski, K Domschke, EJC de Geus, SP Hamilton, C Hayward, I Hickie, JJ Hottenga, M Ising, I Jones, LA Jones, Z Kutalik, S Lucae, NG Martin, Y Milaneschi, B Mueller-Myhsok, MJ Owen, S Padmanabhan, BWJH Penninx, G Pistis, DJ Porteous, M Preisig, S Ripke, SI Shyn, PF Sullivan, J Whitfield, NR Wray, Major Depressive Disorder Working Group of the Psychiatric Genomics Consortium, MEGASTROKE consortium, AM McIntosh, IJ Deary, G Breen, CM Lewis

**Author notes:** Corresponding author: Saskia P Hagenaars,; Social, Genetic and Developmental Psychiatry Centre; Institute of Psychiatry, Psychology & Neuroscience; King’s College London; London SE5 8AF; UK.

## Abstract

**Introduction:** It’s imperative to understand the specific and shared aetiologies of major depression and cardio-metabolic disease, as both traits are frequently comorbid and each represents a major burden to society. This study examined whether there is a genetic association between major depression and cardio-metabolic traits and if this association is stratified by age at onset for major depression.

**Methods:** Polygenic risk scores analysis and linkage disequilibrium score regression was performed to examine whether differences in shared genetic aetiology exist between depression case control status (N cases = 40,940, N controls = 67,532), earlier (N = 15,844), and later onset depression (N = 15,800) with body mass index, coronary artery disease, stroke, and type 2 diabetes in eleven data sets from the Psychiatric Genomics Consortium, Generation Scotland, and UK Biobank.

**Results:** All cardio-metabolic polygenic risk scores were associated with depression status. Significant genetic correlations were found between depression and body mass index, coronary artery disease, and type 2 diabetes. Higher polygenic risk for body mass index, coronary artery disease and type 2 diabetes was associated with both early and later onset depression, while higher polygenic risk for stroke was associated with later onset depression only. Significant genetic correlations were found between body mass index and later onset depression, and between coronary artery disease and both early and late onset depression.

**Conclusions:** The phenotypic associations between major depression and cardio-metabolic traits may partly reflect their overlapping genetic aetiology irrespective of the age depression first presents.

## Introduction

Major depressive disorder (MDD) and cardio-metabolic traits are both major causes of morbidity and mortality in high-income countries. Epidemiological studies have shown a well-established association between them(1): MDD increases the risk of cardio-metabolic disease onset and mortality, but cardio-metabolic disease itself can also increase risk of developing MDD. Specifically, a meta-analysis of 124,509 individuals across 21 studies showed that depression is associated with an 80% increased risk for developing coronary artery disease(2). Mezuk et al(3) showed that MDD predicted a 60% increased risk of type 2 diabetes, while type 2 diabetes predicted a 15% increased risk in MDD. MDD is associated with an increased risk of developing stroke (HR 1.45)(4), but meta-analyses have also shown that ~30% of stroke survivors suffered from MDD(5, 6). Risk factors for cardio-metabolic disease, such as obesity, are also linked to depression. Milaneschi et al(7) showed a bidirectional association between depression and obesity, where obesity increases risk for depression and depression increases risk for subsequent obesity. A more detailed review of the comorbidity between depression and cardio-metabolic traits is given in Penninx, Milaneschi (8).

Multiple mechanisms have been proposed to explain the association between MDD and cardio-metabolic diseases, for example biological dysregulation, or an unhealthy lifestyle(1). However, it remains unclear to what extend these mechanisms influence the association, as most studies examining the mechanisms were based on epidemiological observational study designs. The association between MDD and cardio-metabolic disease could also be due to shared genetic factors. Twin studies have shown that genetic factors contribute ~40% of the variation in liability to both MDD(9) and coronary artery disease(10, 11), 72% to type 2 diabetes(12), and 40 - 70% to BMI(13). No twin heritability estimates are available for stroke, as such studies have very limited numbers of twins. The most recent published GWAS for major depression from the Psychiatric Genomics Consortium, including 135,458 cases and 344,901 controls, identified 44 loci that were significantly associated with major depression, highlighting the highly polygenic nature of major depression(14). The heritability for major depression based on single nucleotide polymorphisms (h^2^_SNP_) was estimated to be 8.7% on the liability scale, based on a lifetime risk of 0.15. Similarly, recent efforts to identify common variants associated with cardio-metabolic disease have shown these traits to be highly polygenic(15–18).

Several studies have shown genetic overlap between MDD and cardio-metabolic traits, in particular with coronary artery disease(14). Findings for other cardio-metabolic traits have been inconsistent. Previous studies have identified genetic overlap between MDD and BMI using polygenic risk scores(19), but not based on genetic correlations(20). However, more recent GWAS studies now have the power to detect a genetic correlation between MDD and BMI(14). Studies using twin data also showed that the phenotypic association between MDD and type 2 diabetes was partly due to genetic effects(21). This finding has been replicated using a polygenic risk score approach(22), but not using genetic correlations(22, 23). The Brainstorm Consortium did not find a significant genetic correlation between MDD and stroke(24), while Wassentheil-Smoller et al(25) showed that higher polygenic risk for MDD was associated with increased risk for stroke, in particular small vessel disease.

The inconsistency in results is likely due to differences in methodological approaches or in summary statistics, but could also be due to the heterogeneity of MDD. MDD onset can occur at any stage of life, but the factors associated with MDD are often age specific or age restricted(26). Increased genetic risk for major depression is associated with earlier AAO compared to later AAO(14). Earlier AAO MDD has a higher heritability and is associated with increased risk for MDD in relatives. On the other hand, vascular disease and its risk factors are linked to a later age at onset (AAO) for MDD(27–29). A large study of Swedish twins showed that a later AAO for MDD in one twin was associated with a higher risk for vascular disease in the other twin(30). To date, no molecular genetic studies have examined the association between late onset MDD and cardio-metabolic traits and in the current study we have more power to replicate and further investigate this association leveraging both summary statistics and common genetic variant information.

The main aim of the present study is to examine the genetic association between MDD and cardio-metabolic traits using polygenic risk scores and genetic correlations. Secondly, we will examine the association stratified by AAO for MDD to test whether a higher genetic predisposition for cardio-metabolic traits is associated with a later AAO for MDD.

## Methods and Materials

### Samples

This study was performed using data from the Psychiatric Genomics Consortium (PGC) MDD working group (PGC-MDD), Generation Scotland: The Scottish Family Health Study (GS:SFHS), and UK Biobank.

#### PGC

Full details of the studies that form the PGC-MDD have previously been published(14). In summary, a subset of eleven studies from the full PGC-MDD analysis were included in the current study, based on the availability of AAO for MDD. All cases were required to have a lifetime diagnosis of MDD based on international consensus criteria (DSM-IV, ICD-9, or ICD-10)(31–33). This was ascertained using structured diagnostic instruments from direct interview by trained interviewers or clinician administered checklists. In most studies (10/11), controls were randomly selected from the general population and were screened for absence of lifetime MDD. This led to a total of 9,518 cases and 11,557 controls with genotype data and AAO information.

#### GS:SFHS

GS:SFHS is a family-based study consisting of 23,690 participants recruited from the population via general medical practices across Scotland. Sample characteristics and recruitment protocols have been described elsewhere(34, 35). In summary, MDD diagnosis was based on the Structured Clinical Interview for DSM-IV disorders (SCID)(36). Participants who answered positively to two mental health screening questions were invited to complete the full SCID to ascertain MDD diagnosis. Cases were further refined through NHS linkage. Controls were defined as participants who answered negatively to the two screening questions or participants who did complete the SCID but did not meet criteria for MDD. This resulted in 1947 cases and 4858 controls, based on unrelated individuals.

#### UK Biobank

UK Biobank is a large resource for identifying determinants of diseases in middle aged and older healthy individuals (www.ukbiobank.ac.uk)(37). A total 502,655 community-dwelling participants aged between 37 and 73 years were recruited between 2006 and 2010 in the United Kingdom, and underwent extensive testing including mental health assessments. MDD status in UK Biobank was derived from the online mental health questionnaire as previously described(38, 39). Briefly, MDD cases were defined as individuals meeting lifetime criteria for MDD based on questions from the Composite International Diagnostic Interview. Individuals reporting previous self-reported diagnosis of schizophrenia (or other psychosis) or bipolar disorder were excluded as MDD cases. Controls were defined as individuals who did not have any self-reported diagnosis of mental illness, did not take any anti-depressant medications, had not previously been hospitalised with a mood disorder, and did not meet previously defined criteria for a mood disorder(40). For the current study, this led to 29,475 cases and 51,243 controls.

### Age at onset (AAO)

AAO was defined as follows, based on previous work by Power et al(26) developed to account for the substantial by-study heterogeneity in the measure. Heterogeneity in AAO within the PGC MDD cohorts has been extensively investigated. Responses depend on the specific setting in which AAO is asked, and may reflect age at first symptoms, first visit to general practitioner or first diagnosis)(26). Using cut offs for AAO (e.g. onset under 30) does not capture the variance in this measure, and we therefore followed Power et al(26) to use the within study distribution to define early and later onset depression. This approach assumes that all cases were recruited from the same age at onset distribution with differences due to study-specific parameters. Cases reporting AAO older than the recorded age at interview were excluded from each study. Within each study, cases were ordered by AAO and divided into equal octiles (O1 – O8). The first three octiles (O1 – O3) were combined into the early AAO group, the last three octiles (O6 – O8) into the late AAO group. Splitting into equal octiles can result in individuals with the same AAO being arbitrarily placed in different octiles. To address this, cases in O4 with the same AAO as the maximum AAO in O3 were assigned to the early AAO group. Similarly, cases in O5 with the same AAO as the minimum AAO in O6 were assigned to the late AAO group. This led to a total of 15,844 MDD cases with early AAO and 15,800 MDD cases with late AAO, and 67,532 controls.

### Genotyping and quality control

Genotyping procedures have been described in the original analysis for each study(14, 41, 42). All analysis were based on individuals from European ancestry only.

PGC cohorts were genotyped following their local protocols. Individual genotype data for all PGC cohorts was processed using the PGC “Ricopili” pipeline for standardised quality control, imputation and analysis (https://github.com/Nealelab/ricopili)(43). Further details on the default parameters used for the current study can be found in the Supplementary material.

For GS:SFHS, genotyping and imputation procedures have previously been published by Nagy et al(41), a brief summary can be found in the Supplementary Material. Monogenic variants and variants with low imputation quality (INFO < 0.4) were removed.

Two highly-overlapping custom genotyping arrays (UK BiLEVE Axiom array and UK Biobank Axiom array, ~800,000 markers) were used to genotype all UK Biobank participants (N = 487,410)(42). Detailed QC procedures are described in the Supplementary Material.

### Statistical analysis

#### Polygenic risk scores

Six polygenic risk scores (PRS) were calculated based on GWAS summary statistics for body mass index (BMI), coronary artery disease (CAD)(16), stroke and two stroke subtypes(15), and type 2 diabetes (T2D)(18) based on external GWAS p-value < 0.5. Table 1 provides further detail on the GWAS summary statistics.

**Table 1.** Information about GWAS summary statistics, based on European individuals only, to create polygenic risk scores in PGC-MDD, GS:SFHS, and UK Biobank.

| Trait | Origin | N cases/N controls | SNPs with<br>p-value <<br>$5*10^{-8}$ |
| --- | --- | --- | --- |
| Body mass index | UK Biobank | 291,683 individuals | 27,164 |
| Coronary artery disease | CARDIoGRAM(16) | 60,801 cases – 123,504<br>controls | 2,213 |
| Stroke: all | MEGASTROKE(15) | 40,585 cases – 406,111<br>controls | 371 |
| Stroke: ischaemic | MEGASTROKE(15) | 34,217 cases – 406,111<br>controls | 307 |
| Stroke: small vessel<br>disease | MEGASTROKE(15) | 5,386 cases – 406,111<br>controls | 0 |
| Type 2 diabetes | DIAGRAM(18) | 26,676 cases – 132,532<br>controls | 1,788 |

PRS were calculated in all genotyped participants in each study using PRSice v2 (https://github.com/choishingwan/PRSice)(45). Prior to creating the scores, clumping was used to obtain SNPs in linkage disequilibrium with an r2 < 0.25 within a 250kb window. Due to the large number of overlapping samples between published BMI GWAS(13) and the PGC studies, a new GWAS was performed in the UK Biobank. The analysis used BGENIE v1.2 (42)on 291,684 individuals in UK Biobank that did not complete the Mental Health Questionnaire and were therefore independent of prediction samples, as MDD case control status was defined from the Mental Health Questionnaire. The Manhattan and QQ plot of the BMI GWAS can be found in the Supplementary Figure 2.

Logistic regression was used to test the associations between the five PRS and four different MDD case-control sets (all MDD cases vs. control subjects, early AAO cases vs. control subjects, late AAO cases vs. control subjects, and late AAO cases vs. early AAO cases). These analyses test our two hypotheses; comparing controls and MDD cases tests the association of genetic risk for cardiometabolic traits with MDD, and whether this association differs by AAO. In contrast, comparing late and early AAO cases tests the association of genetic risk for cardiometabolic traits with AAO itself. Analyses were performed separately for each study, adjusting for relevant covariates in each study (PGC & GS:SFHS: 5 genetic principal components for population stratification; UK Biobank: 6 genetic principal components for population stratification, assessment centre, and genotyping batch). All PRS were standardised with each study across samples. Meta-analyses of the results across studies for each PRS – MDD combination were then conducted to synthesize the findings for maximum statistical power and to check for heterogeneity. Fixed-effects models were used in which the standardized regression coefficients were weighted by the inverse of their squared standard error. We tested for the presence of between-study heterogeneity using Cochran’s Q. We corrected for multiple testing across all 24 meta-analysis models (4 phenotypes × 6 PRS) using the Benjamini Hochberg false discovery rate method(46), using a critical p-value of 0.0026, which means that all p-values equal to or below the critical p-value are considered significant.

#### GWAS meta-analysis and genetic correlations

Genome wide association analyses for three AAO-stratified MDD case-control subsets were performed within each study, with adjustments for population stratification. Study-specific covariates—for example, site or familial relationships—were also fitted as required (see Supplementary Material). The GWAS for GS:SFHS included all individuals (compared to unrelated individuals in the PRS analysis) to maximise power. Quality control of the study-level summary statistics was performed using the EasyQC software(47), which implemented the exclusion of SNPs with imputation quality <0.8 and minor allele count <25. P-value based meta-analyses, with genomic control, were then performed for each of the three outcome measures, using the METAL package(48). SNPs with a combined sample size of less than 1000 participants were excluded.

Genetic correlations between the cardio-metabolic traits and the four MDD summary statistics were calculated using Linkage Disequilibrium score regression (LDSC) (49)using the default HapMap LD reference. To maximise power (and given genetic correlation analyses in LDSC are robust to sample overlap), the largest available summary statistics were used. These included the previously described summary statistics for coronary artery disease(16), stroke(15), and type 2 diabetes(18), as well as summary statistics for major depression from the PGC-MDD and 23andMe GWAS, including UK Biobank(14), and the most recent BMI GWAS, including both UK Biobank and the GIANT consortium(17). All results were corrected for multiple testing using the Benjamini Hochberg false discovery rate method (critical P-value = 0.0064). Differences between genetic correlations were assessed using block-jackknife (Supplementary Material)(50–52).

## Results

### Summary of AAO

The total sample consisted of 40,940 cases, including 15,844 cases classified as early AAO, 15,800 cases as late AAO, 6296 general cases not included in AAO-specific analysis, and 67,532 controls. The mean AAO (SD) across samples was 32.36 (15.22) years. The mean AAO per study ranged from 21.22 years to 37.23 years, with the German PGC-MDD samples, GS:SFHS and UK Biobank having a higher mean AAO. Figure 1 shows the distribution of AAO and the stratification in the early and late AAO groups in each of the studies. Supplementary Figure 1 shows the cumulative distribution of AAO in each study.

**Figure 1.**
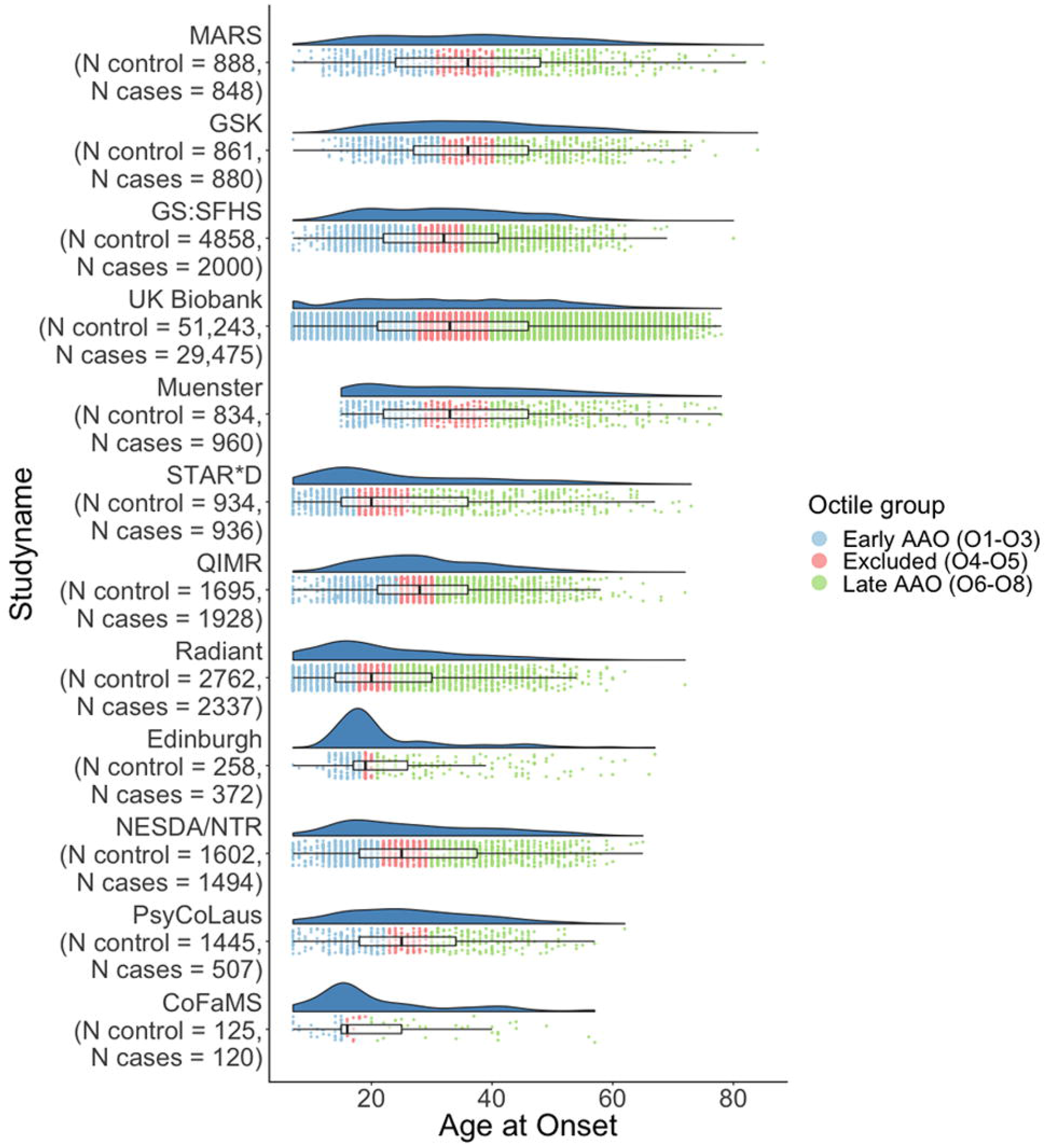
Distribution of Age at onset for MDD per sample. Scatterplot represents the stratification of the early and late AAO groups within each sample.

**Figure 2.**
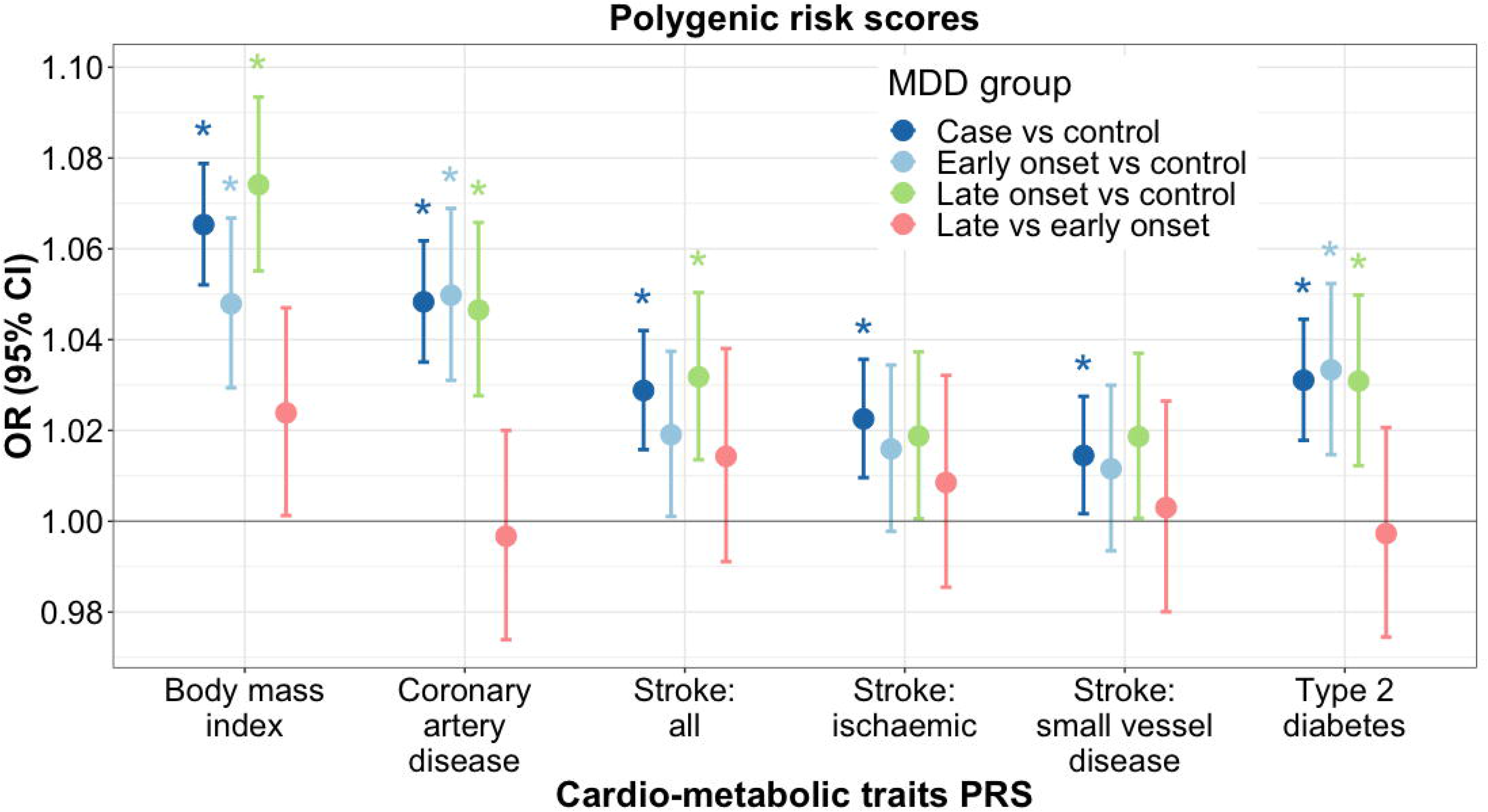
Meta-analysis of associations between cardio-metabolic PRS and stratified MDD outcomes. OR, Odds Ratio; 95% CI, 95% confidence interval; PRS, polygenic risk score.

### Polygenic risk analysis of cardio-metabolic disease

The meta-analyses across studies showed that PRS for BMI, coronary artery disease, all stroke, ischaemic stroke, small vessel disease, and type 2 diabetes were significantly associated with MDD case control status (Figure 2, Supplementary Table 2). Study specific results can be found in Supplementary Table 3.

When stratifying by AAO and comparing with controls, higher PRS for BMI was more strongly associated with late AAO (OR = 1.07, 95% CI = 1.06 – 1.09, P = 4.10 × 10^−15^) than early AAO (OR =1.05, 95% CI = 1.03 – 1.07, P = 2.60 × 10^−7^), however the difference between them was not significant (Figure 2). No significant difference was found between BMI PRS and late vs early AAO within MDD cases (OR = 1.02, 95% CI = 1.00 – 1.05, P = 0.04). Higher PRS for CAD was associated with both late (OR = 1.05, 95% CI = 1.03 – 1.07, P = 8.66 × 10^−7^) and early AAO (OR = 1.05, 95% CI = 1.03 – 1.07, P = 1.50 × 10^−7^) compared to controls, while no significant difference was found between late and early AAO in MDD cases only (OR = 1.00, 95% CI = 0.97 – 1.02, P = 0.776). Higher PRS for all stroke was only associated with late AAO compared to controls (OR = 1.03, 95% CI = 1.01 – 1.05, P = 0.0006). Higher PRS for type 2 diabetes was associated with both late (OR = 1.03, 95% CI = 1.01 – 1.05, P = 0.001) and early (OR = 1.03, 95% CI = 1.01 – 1.05, P = 0.0004) AAO compared to controls, the association between late and early AAO in MDD cases only was non-significant (OR = 1.00, 95% CI = 0.97 – 1.02, P = 0.82).

### Genetic correlations with cardio-metabolic traits

Genetic correlations were calculated between MDD and cardio-metabolic traits based on the largest available GWAS (Figure 3, Supplementary Table 4). Significant genetic correlations were identified between MDD and BMI (r_g_ = 0.13, SE = 0.02, p = 3.43 × 10^−11^), coronary artery disease (r_g_ = 0.12, SE = 0.03, p = 8.47 × 10^−6^), and type 2 diabetes (r_g_ = 0.11, SE = 0.03, p = 0.0001). The genetic correlation between ischaemic stroke and MDD was 0.10 (SE = 0.04, p = 0.026), however this correlation did not survive the correction for multiple testing (p <= 0.0064). No significant genetic correlations were identified for all stroke and small vessel disease.

**Figure 3.**
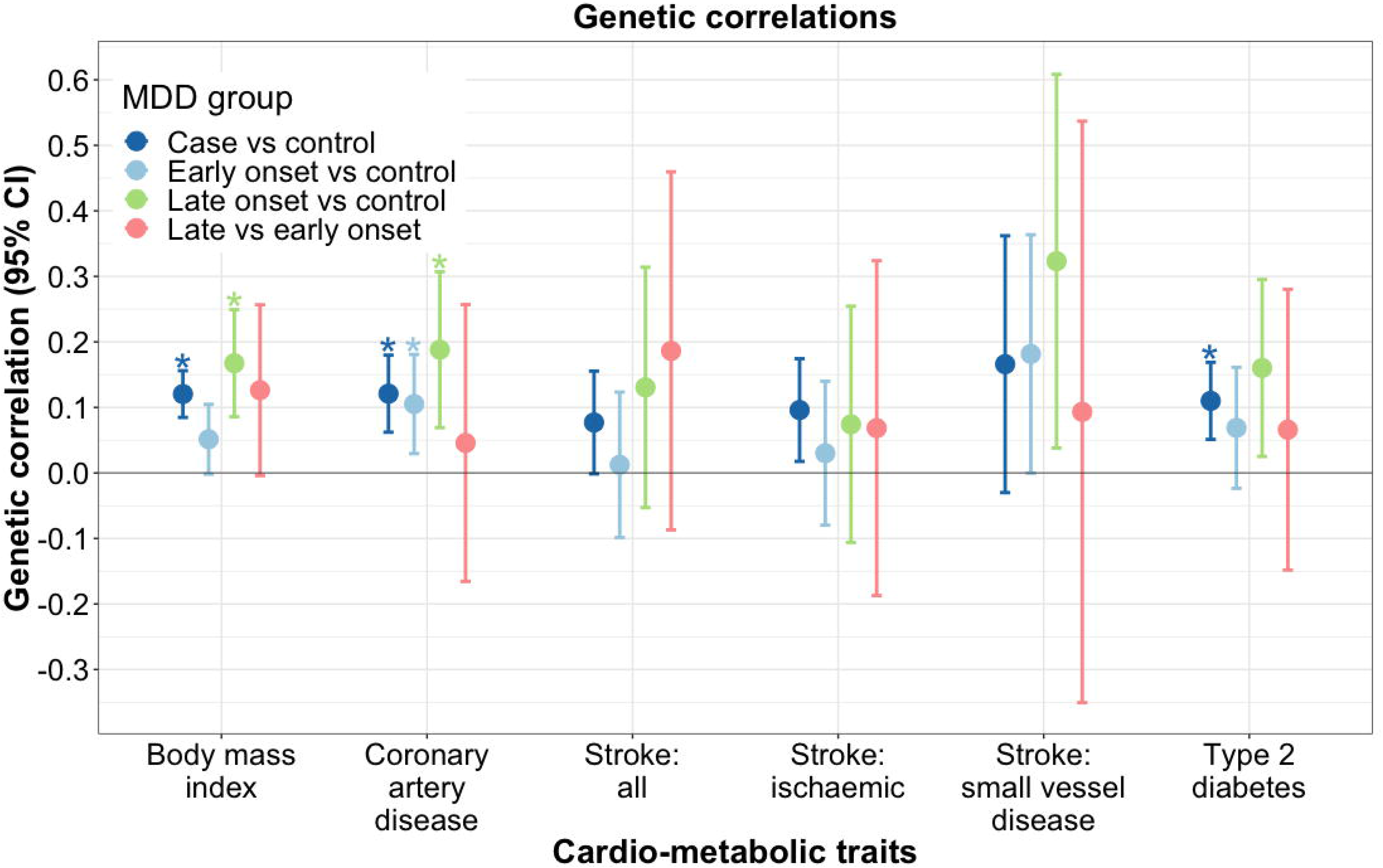
Genetic correlations (SE), derived using LD score regression, between cardio-metabolic traits and stratified MDD outcomes.

In order to calculate genetic correlations between the AAO stratified MDD traits and cardio-metabolic traits, we ran GWAS on the AAO stratified traits. Supplementary Figure 3-5 shows the Manhattan and QQ plots for these traits. Supplementary Table 4 provides further detail on the heritability estimates of the stratified MDD traits. One genome-wide significant variant was identified in the GWAS for early AAO vs controls (rs2789313 on chromosome 10, Z-score = 5.56, P = 2.74 × 10^−8^). This variant is located in the *MALRD1* gene, which is involved in hepatic bile acid metabolism and lipid homeostasis(53). This gene has not previously been associated with depression. Supplementary Figure 3 shows a locus zoom plot of the region on chromosome 10 including rs2789313. Supplementary Tables 5 – 7 provide summary statistics on all suggestive (P < 1 × 10^−5^) variants for the stratified GWAS.

The results for the stratified AAO traits showed a significant genetic correlation between BMI and late AAO vs controls (rg = 0.19, SE = 0.04, p = 2.42 × 10^−5^), no significant genetic correlations were identified for either early AAO vs controls (rg = 0.05, SE = 0.03, p = 0.059) or late vs early AAO (rg = 0.11, SE = 0.05, p = 0.037). A stronger genetic correlation was found between late AAO and CAD (rg = 0.19, SE = 0.06, p = 0.002) compared to early AAO and CAD (rg = 0.11, SE = 0.04, p = 0.006), however the difference between the two genetic correlations was non-significant (block jackknife test for comparison of r_g_ p = 0.227). No significant genetic correlations were found between any of the AAO stratified MDD outcomes and all stroke, ischaemic stroke, small vessel disease, or type 2 diabetes (Figure 3, Supplementary Table 4).

## Discussion

This study explored whether the association between MDD and cardio-metabolic traits is partly due to genetic factors. Using data from publicly available GWAS and from the PGC, UK Biobank, and Generation Scotland, we showed significant genetic overlap between MDD and BMI, coronary artery disease, stroke, and type 2 diabetes from polygenic risk scores and genetic correlations.

Coronary artery disease and type 2 diabetes showed similar patterns of association. Both were associated with MDD in general and with late and early AAO, however the association with late vs early AAO was null. This indicates that both traits are associated with MDD irrespective of the AAO, and not associated with AAO itself. The current study is based on an extension of the strategies applied by Power et al. (26), who showed significant associations between polygenic risk for coronary artery disease and MDD, as well as with early and late onset MDD compared to controls. The results in the current study are based on a larger sample and more recent effect estimates for coronary artery disease, and have therefore greater power to detect an effect.

This is the first study to show genetic overlap between MDD and type 2 diabetes using both polygenic risk scores and genetic correlations. Twin studies have previously shown genetic overlap between MDD and type 2 diabetes, but evidence from genetic studies has been weaker. The current study, however, does show evidence for a shared genetic aetiology between MDD and type 2 diabetes based on both polygenic risk scores and genetic correlations (21–23).

The polygenic risk score analysis showed that individuals with a stronger genetic liability for stroke (and its subtypes) were more likely to suffer from MDD. Clinical and neuroimaging studies have shown that stroke could predispose the development of MDD by disrupting frontostriatal circuits in the brain involved in mood regulation(54–56), specifically in older individuals with MDD or those with a later AAO for MDD. We did not find a significant association between genetic risk for stroke and AAO. This could be due to a lack of power, as the stroke outcomes had low heritability estimates (observed h^2^ ~1%, Supplementary Table 4B) and the sample size of the stratified MDD outcomes is smaller than the overall MDD case-control comparison.

When stratifying by AAO, higher polygenic risk for BMI was more strongly associated with late AAO than early AAO compared to controls, while a significant genetic correlation was only found between late AAO vs control and BMI. A causal association using Mendelian randomization has previously been identified between BMI and depression, showing a 1.12 fold increase in depression for each SD increase in BMI(14). Similarly, Vogelzangs et al(57) have shown that over a five year period both overall and abdominal obesity were associated with an increased risk for MDD onset in men. Although the differences observed in this study are not significant following multiple testing correction, in the context of previous findings, they tentatively suggest that the aetiological processes underlying later onset MDD are linked to vascular pathology and its risk factors such as obesity.

LD score regression produced fewer significant associations than the polygenic risk score analysis. Genetic correlations using LD score regression are based on summary statistics only and could therefore be less powerful than a polygenic risk score approach based on raw genotype data. However, the direction of effect was the same across both analysis approaches and significant genetic correlations were corroborated by significant polygenic risk score associations.

The current study has a number of limitations. The measures for AAO for MDD rely on self-report and are assessed differently across cohorts. We addressed this by stratifying AAO into octiles relative to the mean in each study, therefore assuming that each study recruited AAO from the same distribution with differences between studies due to differences in ascertainment measure. It should be noted that the mean AAO for late onset cases was 47.85 years, which is below what is generally considered to be late onset or geriatric depression (onset > 60 years). More pronounced overlap might exist between late onset MDD and cardio-metabolic traits when focussing on cases with a later AAO, however this study only had a small number of these included. However, this study does show that the effect of genetic risk for somatic conditions on depression is not limited to late onset or geriatric depression. By standardising the PRS in all models we have possibly introduced bias into our results. When the sample prevalence of a trait is not equal to the population prevalence, the mean of the PRS will not represent the mean of the PRS in the population, and thus introducing bias which could lead to inflated effect estimates.

Future studies could focus on the effects of medication or disease status for the cardio-metabolic traits, as the current study was unable to adjust for this due to lack of information on these variables. Pathway analysis might provide further insight into the biology underlying the association between cardio-metabolic traits and MDD. In order to further dissect the comorbidity between MDD and cardio-metabolic disease, it is imperative to have cleaner phenotypes, in particular with regards to AAO. Biobanks with electronic health records might aid in providing improved phenotyping, but this still does not provide the same information as one gets from clinical research studies.

In summary, this study showed genetic overlap between MDD and cardio-metabolic traits based on polygenic risk scores and genetic correlations. These associations were largely irrespective of AAO for MDD, in particular for coronary artery disease and type 2 diabetes. The association with BMI showed some evidence for a stronger link with later onset MDD, however this finding needs to be replicated. Cardio-metabolic traits and its risk factors are to open to preventative strategies, therefore further understanding of the shared genetic aetiology between cardio-metabolic traits and MDD could play an important role in the prevention or management of MDD.

## Acknowledgements

SPH is funded by the Medical Research Council (MR/S0151132). CML is funding by the Medical Research Council (N015746/1). This study presents independent research supported by the National Institute for Health Research (NIHR) Maudsley Biomedical Research Centre at South London and Maudsley NHS Foundation Trust and King’s College London. The views expressed are those of the author(s) and not necessarily those of the NHS, NIHR, Department of Health or King’s College London. We thank participants and scientists involved in making the UK Biobank resource available (http://www.ukbiobank.ac.uk/). UK Biobank data used in this study were obtained under approved application 16577. We are deeply indebted to the investigators who comprise the PGC, and to the hundreds of thousands of subjects who have shared their life experiences with PGC investigators. The PGC has received major funding from the US National Institute of Mental Health and the US National Institute of Drug Abuse (U01 MH109528 and U01 MH1095320). Acknowledgements and funding sources for each of the primary studies in the PGC are contained in supplementary materials. Statistical analyses were carried out on the NL Genetic Cluster Computer (http://www.geneticcluster.org/) hosted by SURFsara, and the King’s Health Partners High Performance Compute Cluster funded with capital equipment grants from the GSTT Charity (TR130505) and Maudsley Charity (980). We would like to thank the research participants and employees of 23andMe for making this work possible. Generation Scotland received core support from the Chief Scientist Office of the Scottish Government Health Directorates [CZD/16/6] and the Scottish Funding Council [HR03006]. Genotyping of the GS:SFHS samples was carried out by the Genetics Core Laboratory at the Wellcome Trust Clinical Research Facility, Edinburgh, Scotland and was funded by the Medical Research Council UK and the Wellcome Trust (Wellcome Trust Strategic Award ‘STratifying Resilience and Depression Longitudinally’ (STRADL) Reference 104036/Z/14/Z). Ethics approval for the Generation Scotland was given by the NHS Tayside committee on research ethics (reference 15/ES/0040), and all participants provided written informed consent for the use of their data. The MEGASTROKE project received funding from sources specified at http://www.megastroke.org/acknowledgments.html

