## Supplementary_material for "Genetic comorbidity between major depression and cardio-metabolic disease, stratified by age at onset of major depression"

Supplementary Methods

- Details of study-specific GWAS of AAO outcomes	2
- Comparing two genetic correlations using block jackknife and LD Score	3

Supplementary Figures

- Supplementary Figure 1: Cumulative distribution of AAO	5
- Supplementary Figure 2: Manhattan and QQ plot for body mass index	6
- Supplementary Figure 3: Manhattan, QQ, Locuszoom plot for early AAO GWAS	7
- Supplementary Figure 4: Manhattan and QQ plot for late AAO GWAS	8
- Supplementary Figure 3: Manhattan and QQ plot for late vs early AAO GWAS	9

Acknowledgments	10

Funding sources	12

Full author list for the Major Depressive Disorder Working Group of the Psychiatric Genomics Consortium	13

Full author list for the MEGASTROKE Consortium	20

**Supplementary Methods**

**Details of study-specific genotype QC**

For the PGC MDD cohorts, the default parameters for retaining SNPs and subjects were: SNP missingness < 0.05 (before sample removal); subject missingness < 0.02; autosomal heterozygosity deviation (|F_het_|<0.2); SNP missingness < 0.02 (after sample removal); difference in SNP missingness between cases and controls < 0.02; and SNP Hardy-Weinberg equilibrium (P>10^−6^ in controls or P>10^−10^ in cases). Genotype imputation was performed using the pre-phasing/imputation stepwise approach implemented in IMPUTE2 / SHAPEIT, based on the 1000 Genomes Project reference panel (August 2012, 30,069,288 variants, release “v3.macGT1”).

For GS:SFHS, genotyping was performed using the Illumina HumanOmniExpressExome-8 v1.0 and v1.2 Beadchip. The quality control parameters for excluding subjects or variants were: subject or variant call rate < 98%, Hardy-Weinberg equilibrium p value less than 1 × 10^–6^, and ancestry outliers based on the 1000 Genomes Project(41). Imputation was performed using the Haplotype Reference Consortium v1.1.

Individuals who had withdrawn consent for analysis, or where recommended by the UK Biobank core analysis team for unusual levels of missingness or heterozygosity were excluded from all analysis. Furthermore, individuals were also excluded based on a genotype call rate < 98%, if they were related to another individual in the dataset (KING r < 0.044, equivalent to removing up to third-degree relatives inclusive)(43), or if the phenotypic and genotypic gender information was discordant (X-chromosome homozygosity (FX) < 0.9 for phenotypic males, FX > 0.5 for phenotypic females) (35). Removal of relatives was performed using a “greedy” algorithm, which minimises exclusions (for example, by excluding the child in a mother-father-child trio). 4-means clustering on the first two genetic principal components (as provided by UK Biobank) was used to define individuals of White Western European ancestry(44). Only these individuals were taken forward for further analysis. Further principal component analysis was performed on this subset of European individuals using the software flashpca2(45) to obtain genetic principal components for use in downstream analyses. After centralised genotype quality control (to remove genotype errors), the genotype data was imputed to the Haplotype Reference Consortium (HRC) and the UK10K reference panels(44).

**Details of study-specific GWAS of AAO outcomes.**

PGC

GWAS of each of the stratified MDD outcomes were conducted using the postimp-navi module of the Ricopili pipeline, adjusting for 5 principal components. One of the PGC samples (Muenster) has been analysed separately from the original PGC cohorts using the Ricopili pipeline, as this sample was added later.

Generation Scotland

GWAS of each of the stratified MDD outcomes in GS:SFHS were conducted using mixed linear model based association (MLMA) analysis, implemented in GCTA (v1.25.). Two genomic relationship matrices (GRM) were used to account for population structure, to allow inclusion of closely and distantly related individuals. The first GRM included pairwise relationship coefficients for all individuals. The second GRM had off-diagonal elements < 0.05 set to 0. GRMs were created using the mixed linear model with candidate marker excluded (MLMe) approach, where GRMs are calculated excluding SNPs located on the chromosome under analysis. MLMA employs restricted maximum likelihood methods on the linear scale. As such, test statistics (betas and their corresponding standard errors) were transformed to Odds Ratios and their corresponding 95% Confidence Intervals on the liability scale using a Taylor transformation expansion series. Further details have been published previously^1^.

UK Biobank

For UK Biobank, GWAS on the AAO stratified MDD outcomes were conducted using linear regressions on imputed genotype dosages in BGenie v1.2, with residualised phenotypes. Deviance residuals were obtained from logistic regressions in R.3.4.1. Six principal components, genotype batch, and assessment centre were used to control for geographical variation in the data.

**Comparing two genetic correlations using block jackknife and LD Score**

Define four phenotypes: A, B, C, and D. We wished to compare the genetic correlation of A and B to the genetic correlation of C and D. Global estimates of these correlations, denoted $r\left( A,B \right)$ and $r\left( C,D \right)$, can be computed using LD Score. The same software can output jackknife delete values for genetic covariance: $G\left( A,B \right)$, $G\left( C,D \right)$, as well as for heritability: $H\left( A,B \right)$ and $H\left( C,D \right)$. These jackknife delete values are estimated by excluding blocks of values (here, number of blocks n = 200). The n-dimensional vectors $G\left( A,B \right)$, $G\left( C,D \right)$, $H\left( A,B \right)$ and $H\left( C,D \right)$ can be used to generate genetic correlation delete values $R\left( A,B \right)$ and $R\left( C,D \right)$. The difference between the global estimates $r\left( A,B \right)$ and $r\left( C,D \right)$ is $d\left( AB,CD \right)$ and the difference between the vectors $R\left( A,B \right)$ and $R\left( C,D \right)$ is $D\left( AB,CD \right)$. The global genetic correlation difference $d\left( AB,CD \right)$ and the delete values $D\left( AB,CD \right)$ are used to compute jackknife pseudovalues. The ith pseudovalue is:

$$P_{i}\left( AB,CD \right)=n\times d\left( AB,CD \right)-\left( n-1 \right)*D_{i}\left( AB,CD \right)$$

The mean and variance of the jackknife pseudovalues are:

$$m(AB,CD)=\frac{1}{n}\sum_{i=1}^{n} P_{i}(AB,CD)$$

$$v(AB,CD)=\frac{1}{n-1}\sum_{i=1}^{n} {(P_{i}(AB,CD)-m(AB,CD))}^{2}$$

The jackknife estimate of the difference between the two correlations m(AB,CD) can then be compared to test H0 : θ = θ_0_ (where θ_0_ = 0 for no difference between genetic correlations), and a p-value can be derived from the z statistic:

$$z(AB,CD)=\frac{m(AB,CD)-\theta_{0}}{\sqrt{(1/n)\times v(AB,CD)}}$$

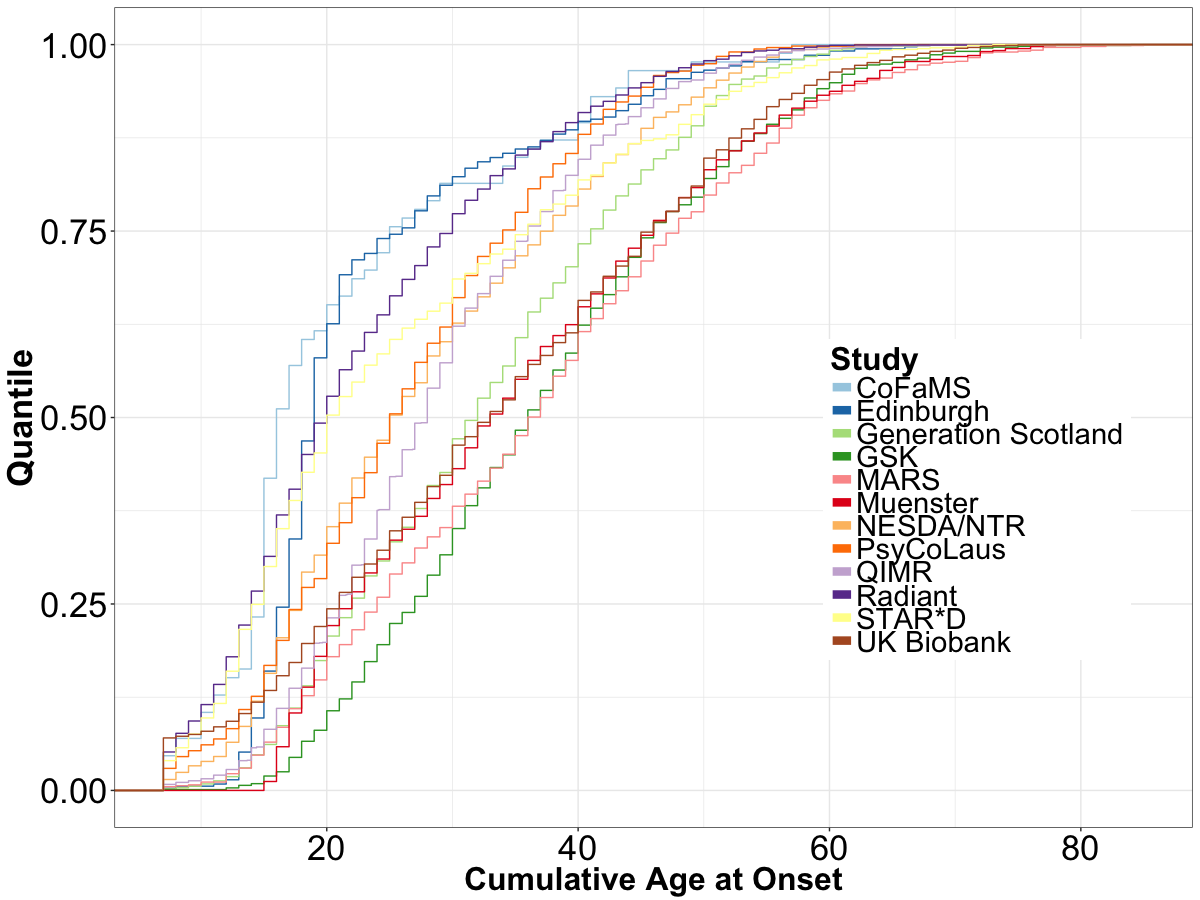

**Supplementary Figure 1**. Cumulative distribution of AAO in all studies.

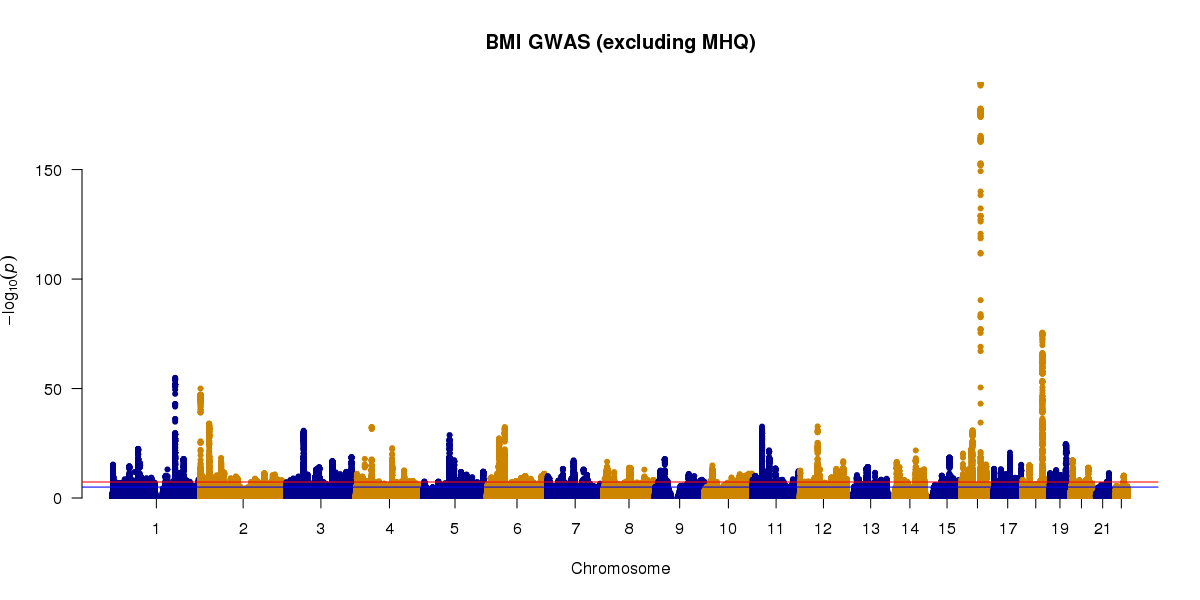

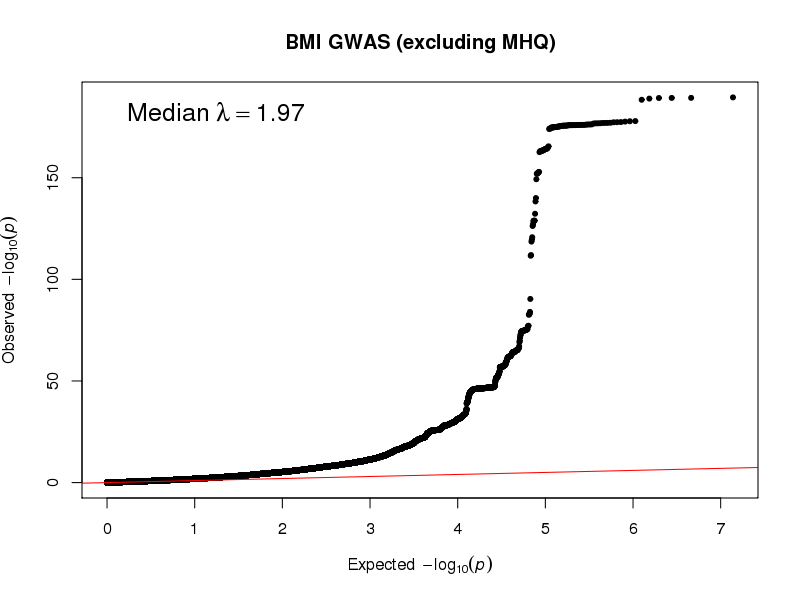

**Supplementary Figure 2.** Manhattan and QQ plot for body mass index in UK Biobank in samples that exclude individuals who completed the Mental health questionnaire.

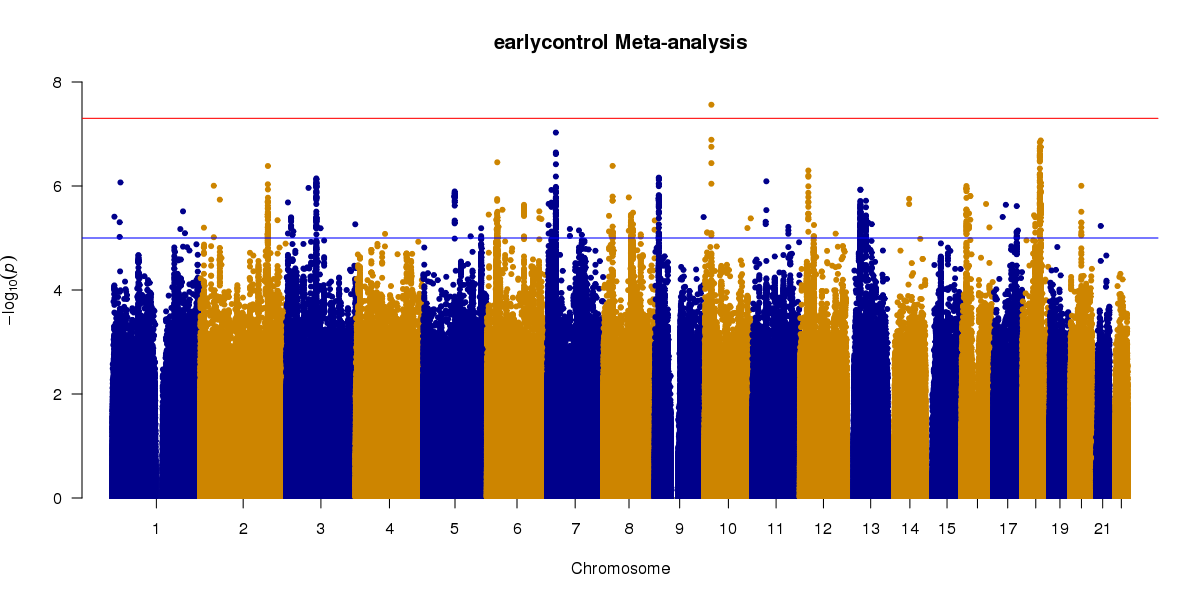

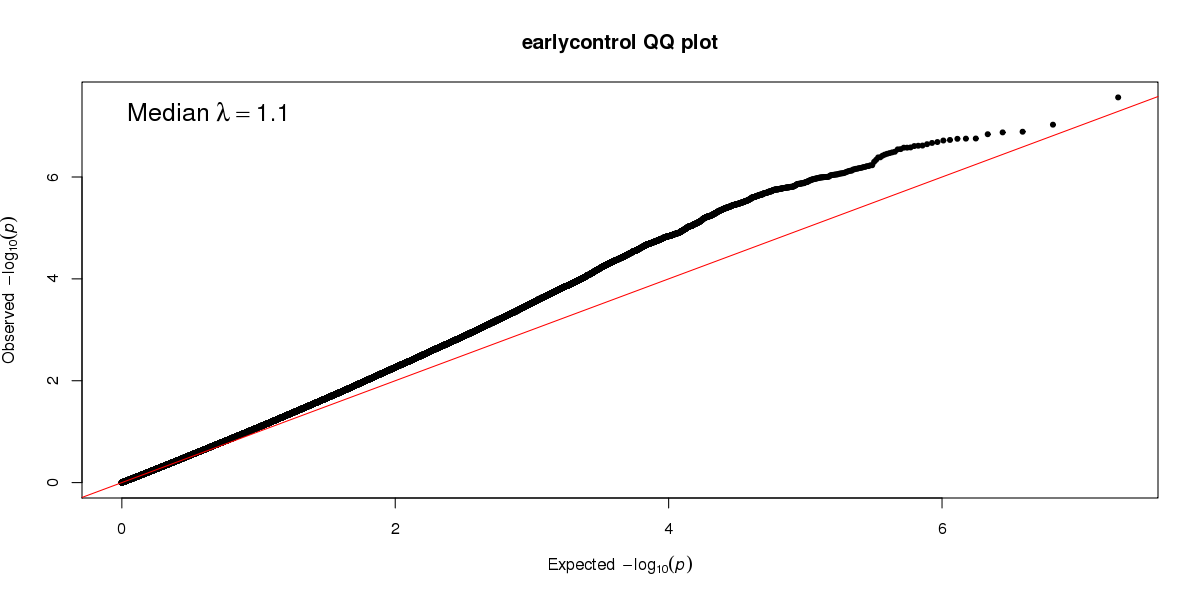

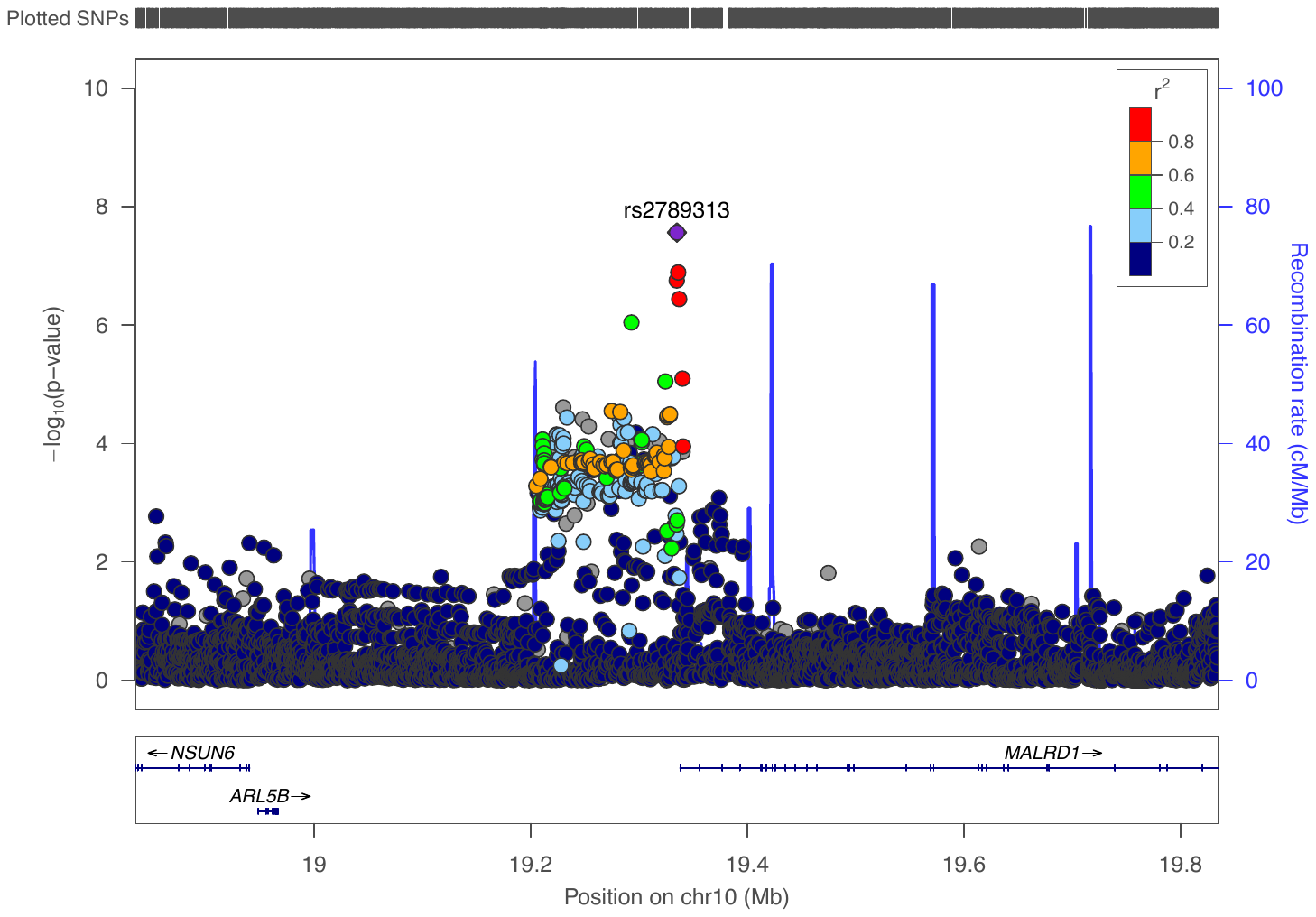

**Supplementary Figure 3.** Manhattan, QQ, and Locus zoom plot for early AAO vs control GWAS in UK Biobank in samples that exclude individuals who completed the Mental health questionnaire.

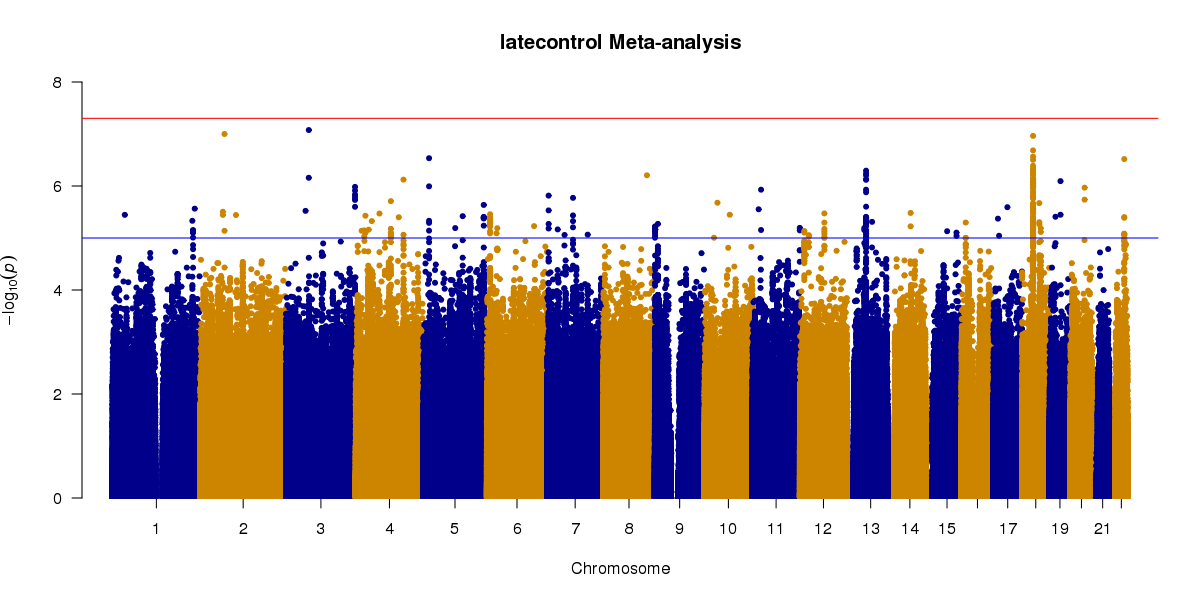

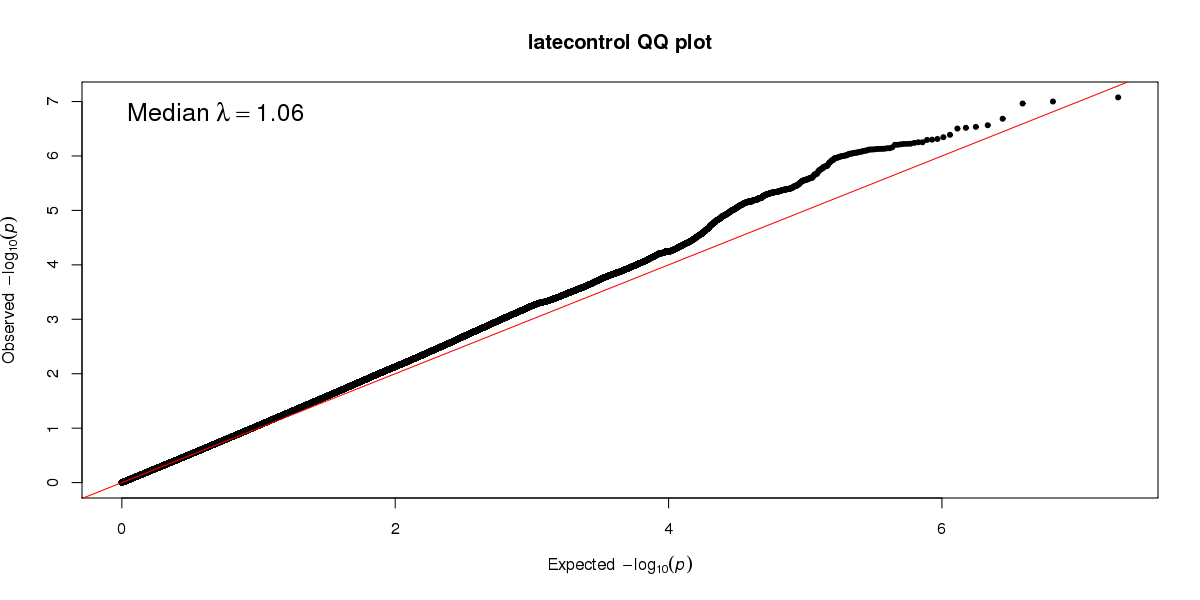

**Supplementary Figure 4.** Manhattan and QQ plot for late AAO vs control GWAS in UK Biobank in samples that exclude individuals who completed the Mental health questionnaire.

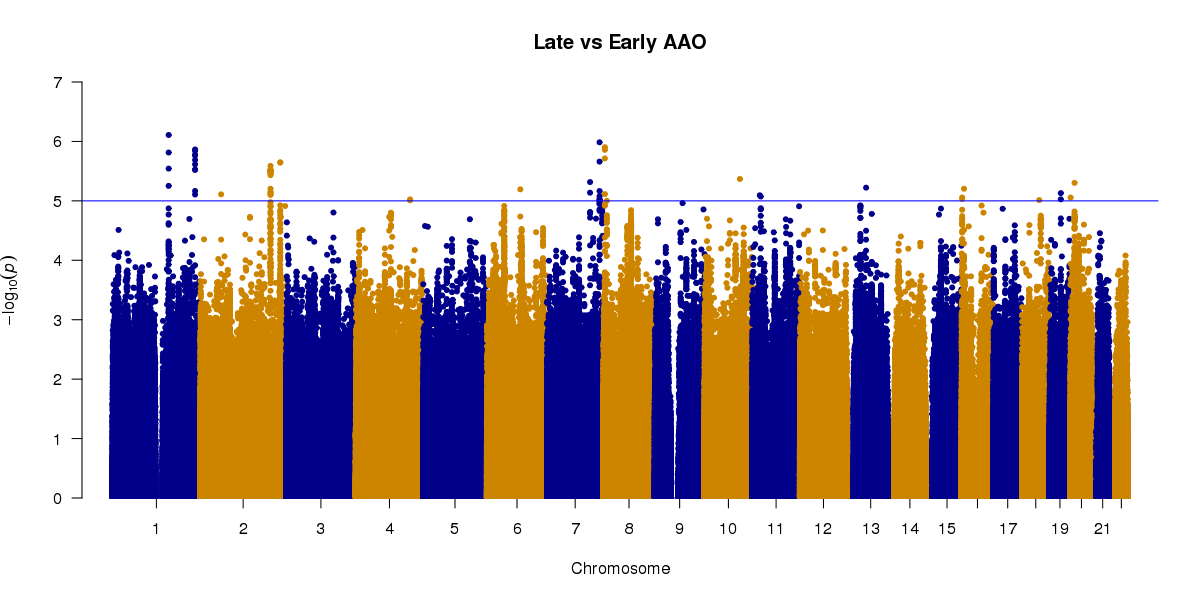

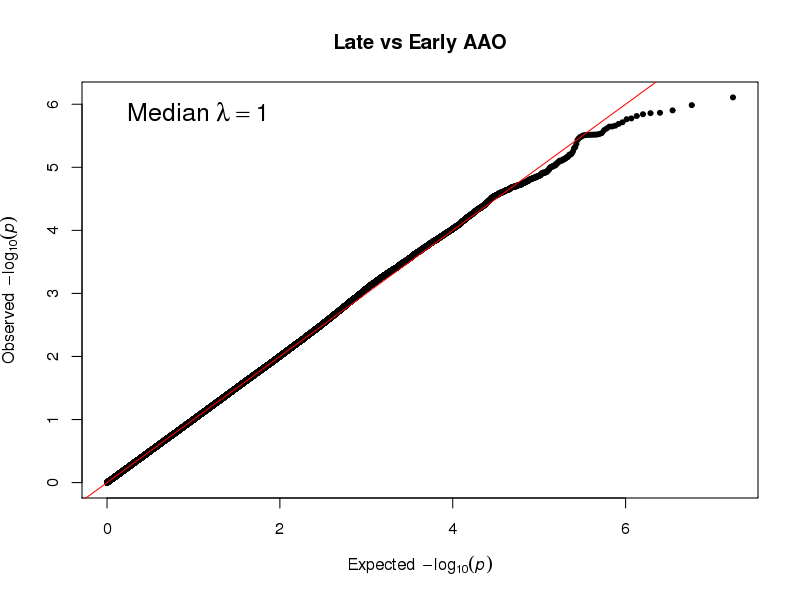

**Supplementary Figure 5.** Manhattan and QQ plot for late AAO vs control GWAS in UK Biobank in samples that exclude individuals who completed the Mental health questionnaire.

**Acknowledgements**

We thank contributors to the primary studies comprising the PGC sample in this paper. **EDINBURGH**: Genotyping was conducted at the Genetics Core Laboratory at the Clinical Research Facility (University of Edinburgh). **GSK_MUNICH**: We thank all participants in the GSK-Munich study. We thank numerous people at GSK and Max-Planck Institute, BKH Augsburg and Klinikum Ingolstadt in Germany who contributed to this project. **MARS**: This work was funded by the Max Planck Society, by the Max Planck Excellence Foundation, and by a grant from the German Federal Ministry for Education and Research (BMBF) in the National Genome Research Network framework (NGFN2 and NGFN-Plus, FKZ 01GS0481), and by the BMBF Program FKZ 01ES0811. We acknowledge all study participants. We thank numerous people at Max-Planck Institute, and all study sites in Germany and Switzerland who contributed to this project. Controls were from the Dortmund Health Study which was supported by the German Migraine & Headache Society, and by unrestricted grants to the University of Münster from Almirall, Astra Zeneca, Berlin Chemie, Boehringer, Boots Health Care, Glaxo-Smith-Kline, Janssen Cilag, McNeil Pharma, MSD Sharp & Dohme, and Pfizer. Blood collection was funded by the Institute of Epidemiology and Social Medicine, University of Münster. Genotyping was supported by the German Ministry of Research and Education (BMBF grant 01ER0816). **NESDA:** The infrastructure for the NESDA study ([www.nesda.nl](http://www.nesda.nl/)) is funded through the Geestkracht program of the Netherlands Organisation for Health Research and Development (ZonMw, grant number 10-000-1002) and financial contributions by participating universities and mental health care organizations (VU University Medical Center, GGZ inGeest, Leiden University Medical Center, Leiden University, GGZ Rivierduinen, University Medical Center Groningen, University of Groningen, Lentis, GGZ Friesland, GGZ Drenthe, Rob Giel Onderzoekscentrum). Genotyping was supported by the Biobanking and Biomolecular Resources Research Infrastructure (BBMRI-NL) and National Institutes of Health (Genetic Association Information Network (GAIN) of the Foundation for the National Institutes of Health, Grand Opportunity grants 1RC2 MH089951 and 1RC2 MH089995). **NTR**: is supported by NWO(480-15-001/674); BBMRI –NL: Biobanking and Biomolecular Resources Research Infrastructure (184.021.007 and BBMRI-NL2.0 184.033.111); Genotyping was supported BBMRI-NL, the Avera Institute, Sioux Falls (USA) and National Institutes of Health (Genetic Association Information Network (GAIN) of the Foundation for the National Institutes of Health, Grand Opportunity grants 1RC2 MH089951 and 1RC2 MH089995). **PsyColaus**: PsyCoLaus/CoLaus received additional support from research grants from GlaxoSmithKline and the Faculty of Biology and Medicine of Lausanne. **QIMR**: We thank the twins and their families for their willing participation in our studies. **RADIANT**: This report represents independent research funded by the National Institute for Health Research (NIHR) Biomedical Research Centre at South London and Maudsley NHS Foundation Trust, and King’s College London. The views expressed are those of the authors and not necessarily those of the NHS, the NIHR, or the Department of Health. **STAR*D**: The authors appreciate the efforts of the STAR*D investigator team for acquiring, compiling, and sharing the STAR*D clinical data set.

Funding support for the Genome-Wide Association of Schizophrenia Study was provided by the National Institute of Mental Health (R01 MH67257, R01 MH59588, R01 MH59571, R01 MH59565, R01 MH59587, R01 MH60870, R01 MH59566, R01 MH59586, R01 MH61675, R01 MH60879, R01 MH81800, U01 MH46276, U01 MH46289 U01 MH46318, U01 MH79469, and U01 MH79470) and the genotyping of samples was provided through the Genetic Association Information Network (GAIN). The datasets used for the analyses described in this manuscript were obtained from the database of Genotypes and Phenotypes (dbGaP) found at <http://www.ncbi.nlm.nih.gov/gap> through dbGaP accession number phs000021.v3.p2. Samples and associated phenotype data for the Genome-Wide Association of Schizophrenia Study were provided by the Molecular Genetics of Schizophrenia Collaboration (PI: Pablo V. Gejman, Evanston Northwestern Healthcare (ENH) and Northwestern University, Evanston, IL, USA).

**Funding sources**

The table below lists the funding that supported the primary studies analysed in this paper.

| **Study** | **Lead investigator** | **Award number** | **Funder** | **Country** |
| --- | --- | --- | --- | --- |
| PGC | PF Sullivan | U01 MH109528 | NIMH | USA |
| PGC | A Agrawal | U01 MH109532 | NIDA | USA |
| PGC | D Posthuma | 480-05-003 | Netherlands Scientific Organization | Netherlands |
| PGC | D Posthuma | – | Dutch Brain Foundation and the VU University Amsterdam | Netherlands |
| CoFaMS - Adelaide | BT Baune | APP1060524 | NHMRC | Australia |
| Generation Scotland | AM McIntosh | 104036/Z/14/Z | Wellcome Trust | UK |
| Münster MDD cohort | BT Baune | N Health-F2-2008-222963 | European Union | Germany |
| Münster MDD cohort | TG Schulze | 01ZX1314K | BMBF Integrament | Germany |
| Münster MDD cohort | TG Schulze |  | Dr. Lisa Oehler Foundation | Germany |
| Münster MDD cohort | TG Schulze | SCHU 1603/5-1 | German Research Foundation (DFG) | Germany |
| Münster MDD cohort | U Dannlowski | FOR2107 DA1151/5-1; SFB- TRR58, Project C09 | German Research Foundation (DFG) | Germany |
| Münster MDD cohort | U Dannlowski | Dan3/012/17 | Interdisciplinary Center for Clinical Research, Medical Faculty of University of Münster | Germany |
| Münster MDD cohort | V Arolt | N Health-F2-2008-222963 | European Union | Germany |
| NESDA | BWJH Penninx | ZonMW Geestkracht grant | N.W.O. | Netherlands |
| NESDA | BJWH Penninx | 1RC2 MH089951; 1RC2 MH089995 | NIH | USA |
| NTR | DI Boomsma | 1RC2 MH089951; 1RC2 MH089995 | NIH | Netherlands |
| PsyColaus | M Preisig | 3200B0–105993, 3200B0-118308, 33CSCO-122661, 33CS30-139468, 33CS30-148401 | Swiss National Science Foundation | Switzerland |
| QIMR | NG Martin | 941177, 971232, 3399450 and 443011 | National Health and Medical Research Council | Australia |
| QIMR | AC Heath | AA07535, AA07728, andAA10249 | NIAAA | USA |
| RADIANT | C Lewis, G Breen | G0701420 | MRC | UK |
| RADIANT | G Breen | G0901245 | MRC | UK |
| RADIANT | G Breen | U01 MH109528 | NIMH | UK |
| STAR*D | SP Hamilton | R01 MH-072802 | NIMH | USA |
| UK Biobank | G Breen |  | NIHR | UK |

**Full author list for the Major Depressive Disorder Working Group of the Psychiatric Genomics Consortium**

Naomi R Wray* 1, 2

Stephan Ripke* 3, 4, 5

Manuel Mattheisen* 6, 7, 8, 9

Maciej Trzaskowski* 1

Enda M Byrne 1

Abdel Abdellaoui 10

Mark J Adams 11

Esben Agerbo 9, 12, 13

Tracy M Air 14

Till F M Andlauer 15, 16

Silviu-Alin Bacanu 17

Marie Bækvad-Hansen 9, 18

Aartjan T F Beekman 19

Tim B Bigdeli 17, 20

Elisabeth B Binder 15, 21

Douglas H R Blackwood 11

Julien Bryois 22

Henriette N Buttenschøn 9, 23

Jonas Bybjerg-Grauholm 9, 18

Na Cai 24, 25

Enrique Castelao 26

Jane Hvarregaard Christensen 7, 8, 9

Toni-Kim Clarke 11

Jonathan R I Coleman 27

Lucía Colodro-Conde 28

Baptiste Couvy-Duchesne 2, 29

Nick Craddock 30

Gregory E Crawford 31, 32

Gail Davies 33

Ian J Deary 33

Franziska Degenhardt 34, 35

Eske M Derks 28

Nese Direk 36, 37

Conor V Dolan 10, 118

Erin C Dunn 38, 39, 40

Thalia C Eley 27

Valentina Escott-Price 41

Farnush Farhadi Hassan Kiadeh 42

Hilary K Finucane 43, 44

Jerome C Foo 45

Andreas J Forstner 34, 35, 46, 47

Josef Frank 45

Héléna A Gaspar 27

Michael Gill 48

Fernando S Goes 49

Scott D Gordon 28

Jakob Grove 7, 8, 9, 50

Lynsey S Hall 11, 51

Christine Søholm Hansen 9, 18

Thomas F Hansen 52, 53, 54

Stefan Herms 34, 35, 47

Ian B Hickie 55

Per Hoffmann 34, 35, 47

Georg Homuth 56

Carsten Horn 57

Jouke-Jan Hottenga 10,118

David M Hougaard 9, 18

Marcus Ising 58

Rick Jansen 19

Ian Jones 59

Lisa A Jones 60

Eric Jorgenson 61

James A Knowles 62

Isaac S Kohane 63, 64, 65

Julia Kraft 4

Warren W. Kretzschmar 66

Jesper Krogh 67

Zoltán Kutalik 68, 69

Yihan Li 66

Penelope A Lind 28

Donald J MacIntyre 70, 71

Dean F MacKinnon 49

Robert M Maier 2

Wolfgang Maier 72

Jonathan Marchini 73

Hamdi Mbarek 10, 118

Patrick McGrath 74

Peter McGuffin 27

Sarah E Medland 28

Divya Mehta 2, 75

Christel M Middeldorp 10, 76, 77

Evelin Mihailov 78

Yuri Milaneschi 19

Lili Milani 78

Francis M Mondimore 49

Grant W Montgomery 1

Sara Mostafavi 79, 80

Niamh Mullins 27

Matthias Nauck 81, 82

Bernard Ng 80

Michel G Nivard 10, 118

Dale R Nyholt 83

Paul F O'Reilly 27

Hogni Oskarsson 84

Michael J Owen 59

Jodie N Painter 28

Carsten Bøcker Pedersen 9, 12, 13

Marianne Giørtz Pedersen 9, 12, 13

Roseann E. Peterson 17, 85

Erik Pettersson 22

Wouter J Peyrot 19

Giorgio Pistis 26

Danielle Posthuma 86, 87

Jorge A Quiroz 88

Per Qvist 7, 8, 9

John P Rice 89

Brien P. Riley 17

Margarita Rivera 27, 90

Saira Saeed Mirza 36

Robert Schoevers 91

Eva C Schulte 92, 93

Ling Shen 61

Jianxin Shi 94

Stanley I Shyn 95

Engilbert Sigurdsson 96

Grant C B Sinnamon 97

Johannes H Smit 19

Daniel J Smith 98

Hreinn Stefansson 99

Stacy Steinberg 99

Fabian Streit 45

Jana Strohmaier 45

Katherine E Tansey 100

Henning Teismann 101

Alexander Teumer 102

Wesley Thompson 9, 53, 103, 104

Pippa A Thomson 105

Thorgeir E Thorgeirsson 99

Matthew Traylor 106

Jens Treutlein 45

Vassily Trubetskoy 4

André G Uitterlinden 107

Daniel Umbricht 108

Sandra Van der Auwera 109

Albert M van Hemert 110

Alexander Viktorin 22

Peter M Visscher 1, 2

Yunpeng Wang 9, 53, 104

Bradley T. Webb 111

Shantel Marie Weinsheimer 9, 53

Jürgen Wellmann 101

Gonneke Willemsen 10,118

Stephanie H Witt 45

Yang Wu 1

Hualin S Xi 112

Jian Yang 2, 113

Futao Zhang 1

Volker Arolt 114

Bernhard T Baune 115

Klaus Berger 101

Dorret I Boomsma 10,118

Sven Cichon 34, 47, 116, 117

Udo Dannlowski 114

EJC de Geus 10, 118

J Raymond DePaulo 49

Enrico Domenici 119

Katharina Domschke 120

Tõnu Esko 5, 78

Hans J Grabe 109

Steven P Hamilton 121

Caroline Hayward 122

Andrew C Heath 89

Kenneth S Kendler 17

Stefan Kloiber 58, 123, 124

Glyn Lewis 125

Qingqin S Li 126

Susanne Lucae 58

Pamela AF Madden 89

Patrik K Magnusson 22

Nicholas G Martin 28

Andrew M McIntosh 11, 33

Andres Metspalu 78, 127

Ole Mors 9, 128

Preben Bo Mortensen 8, 9, 12, 13

Bertram Müller-Myhsok 15, 16, 129

Merete Nordentoft 9, 130

Markus M Nöthen 34, 35

Michael C O'Donovan 59

Sara A Paciga 131

Nancy L Pedersen 22

Brenda WJH Penninx 19

Roy H Perlis 38, 132

David J Porteous 105

James B Potash 133

Martin Preisig 26

Marcella Rietschel 45

Catherine Schaefer 61

Thomas G Schulze 45, 93, 134, 135, 136

Jordan W Smoller 38, 39, 40

Kari Stefansson 99, 137

Henning Tiemeier 36, 138, 139

Rudolf Uher 140

Henry Völzke 102

Myrna M Weissman 74, 141

Thomas Werge 9, 53, 142

Cathryn M Lewis 27, 143

Douglas F Levinson 144

Gerome Breen 27, 145

Anders D Børglum 7, 8, 9

Patrick F Sullivan 22, 146, 147,

1, Institute for Molecular Bioscience, The University of Queensland, Brisbane, QLD, AU

2, Queensland Brain Institute, The University of Queensland, Brisbane, QLD, AU

3, Analytic and Translational Genetics Unit, Massachusetts General Hospital, Boston, MA, US

4, Department of Psychiatry and Psychotherapy, Universitätsmedizin Berlin Campus Charité Mitte, Berlin, DE

5, Medical and Population Genetics, Broad Institute, Cambridge, MA, US

6, Centre for Psychiatry Research, Department of Clinical Neuroscience, Karolinska Institutet, Stockholm, SE

7, Department of Biomedicine, Aarhus University, Aarhus, DK

8, iSEQ, Centre for Integrative Sequencing, Aarhus University, Aarhus, DK

9, iPSYCH, The Lundbeck Foundation Initiative for Integrative Psychiatric Research, DK

10 Netherlands Twin Register, Dept of Biological Psychology, Vrije Universiteit Amsterdam, Amsterdam, NL

11, Division of Psychiatry, University of Edinburgh, Edinburgh, GB

12, Centre for Integrated Register-based Research, Aarhus University, Aarhus, DK

13, National Centre for Register-Based Research, Aarhus University, Aarhus, DK

14, Discipline of Psychiatry, Adelaide Medical School, University of Adelaide, Adelaide, SA, AU

15, Department of Translational Research in Psychiatry, Max Planck Institute of Psychiatry, Munich, DE

16, Munich Cluster for Systems Neurology (SyNergy), Munich, DE

17, Department of Psychiatry, Virginia Commonwealth University, Richmond, VA, US

18, Center for Neonatal Screening, Department for Congenital Disorders, Statens Serum Institut, Copenhagen, DK

19, Department of Psychiatry, Amsterdam UMC, Vrije Universiteit Amsterdam, NL

20, Virginia Institute for Psychiatric and Behavior Genetics, Richmond, VA, US

21, Department of Psychiatry and Behavioral Sciences, Emory University School of Medicine, Atlanta, GA, US

22, Department of Medical Epidemiology and Biostatistics, Karolinska Institutet, Stockholm, SE

23, NIDO | Danmark, Regional Hospital West Jutland, DK

24, Human Genetics, Wellcome Trust Sanger Institute, Cambridge, GB

25, Statistical genomics and systems genetics, European Bioinformatics Institute (EMBL-EBI), Cambridge, GB

26, Department of Psychiatry, University Hospital of Lausanne, Prilly, Vaud, CH

27, Social Genetic and Developmental Psychiatry Centre, King's College London, London, GB

28, Genetics and Computational Biology, QIMR Berghofer Medical Research Institute, Brisbane, QLD, AU

29, Centre for Advanced Imaging, The University of Queensland, Brisbane, QLD, AU

30, Psychological Medicine, Cardiff University, Cardiff, GB

31, Center for Genomic and Computational Biology, Duke University, Durham, NC, US

32, Department of Pediatrics, Division of Medical Genetics, Duke University, Durham, NC, US

33, Centre for Cognitive Ageing and Cognitive Epidemiology, University of Edinburgh, Edinburgh, GB

34, Institute of Human Genetics, University of Bonn, Bonn, DE

35, Life&Brain Center, Department of Genomics, University of Bonn, Bonn, DE

36, Epidemiology, Erasmus MC, Rotterdam, Zuid-Holland, NL

37, Psychiatry, Dokuz Eylul University School Of Medicine, Izmir, TR

38, Department of Psychiatry, Massachusetts General Hospital, Boston, MA, US

39, Psychiatric and Neurodevelopmental Genetics Unit (PNGU), Massachusetts General Hospital, Boston, MA, US

40, Stanley Center for Psychiatric Research, Broad Institute, Cambridge, MA, US

41, Neuroscience and Mental Health, Cardiff University, Cardiff, GB

42, Bioinformatics, University of British Columbia, Vancouver, BC, CA

43, Department of Epidemiology, Harvard T.H. Chan School of Public Health, Boston, MA, US

44, Department of Mathematics, Massachusetts Institute of Technology, Cambridge, MA, US

45, Department of Genetic Epidemiology in Psychiatry, Central Institute of Mental Health,  Medical Faculty Mannheim, Heidelberg University, Mannheim, Baden-Württemberg, DE

46, Department of Psychiatry (UPK), University of Basel, Basel, CH

47, Human Genomics Research Group, Department of Biomedicine, University of Basel, Basel, CH

48, Department of Psychiatry, Trinity College Dublin, Dublin, IE

49, Psychiatry & Behavioral Sciences, Johns Hopkins University, Baltimore, MD, US

50, Bioinformatics Research Centre, Aarhus University, Aarhus, DK

51, Institute of Genetic Medicine, Newcastle University, Newcastle upon Tyne, GB

52, Danish Headache Centre, Department of Neurology, Rigshospitalet, Glostrup, DK

53, Institute of Biological Psychiatry, Mental Health Center Sct. Hans, Mental Health Services Capital Region of Denmark, Copenhagen, DK

54, iPSYCH, The Lundbeck Foundation Initiative for Psychiatric Research, Copenhagen, DK

55, Brain and Mind Centre, University of Sydney, Sydney, NSW, AU

56, Interfaculty Institute for Genetics and Functional Genomics, Department of Functional Genomics, University Medicine and Ernst Moritz Arndt University Greifswald, Greifswald, Mecklenburg-Vorpommern, DE

57, Roche Pharmaceutical Research and Early Development, Pharmaceutical Sciences, Roche Innovation Center Basel, F. Hoffmann-La Roche Ltd, Basel, CH

58, Max Planck Institute of Psychiatry, Munich, DE

59, MRC Centre for Neuropsychiatric Genetics and Genomics, Cardiff University, Cardiff, GB

60, Department of Psychological Medicine, University of Worcester, Worcester, GB

61, Division of Research, Kaiser Permanente Northern California, Oakland, CA, US

62, Psychiatry & The Behavioral Sciences, University of Southern California, Los Angeles, CA, US

63, Department of Biomedical Informatics, Harvard Medical School, Boston, MA, US

64, Department of Medicine, Brigham and Women's Hospital, Boston, MA, US

65, Informatics Program, Boston Children's Hospital, Boston, MA, US

66, Wellcome Trust Centre for Human Genetics, University of Oxford, Oxford, GB

67, Department of Endocrinology at Herlev University Hospital, University of Copenhagen, Copenhagen, DK

68, Institute of Social and Preventive Medicine (IUMSP), University Hospital of Lausanne, Lausanne, VD, CH

69, Swiss Institute of Bioinformatics, Lausanne, VD, CH

70, Division of Psychiatry, Centre for Clinical Brain Sciences, University of Edinburgh, Edinburgh, GB

71, Mental Health, NHS 24, Glasgow, GB

72, Department of Psychiatry and Psychotherapy, University of Bonn, Bonn, DE

73, Statistics, University of Oxford, Oxford, GB

74, Psychiatry, Columbia University College of Physicians and Surgeons, New York, NY, US

75, School of Psychology and Counseling, Queensland University of Technology, Brisbane, QLD, AU

76, Child and Youth Mental Health Service, Children's Health Queensland Hospital and Health Service, South Brisbane, QLD, AU

77, Child Health Research Centre, University of Queensland, Brisbane, QLD, AU

78, Estonian Genome Center, University of Tartu, Tartu, EE

79, Medical Genetics, University of British Columbia, Vancouver, BC, CA

80, Statistics, University of British Columbia, Vancouver, BC, CA

81, DZHK (German Centre for Cardiovascular Research), Partner Site Greifswald, University Medicine, University Medicine Greifswald, Greifswald, Mecklenburg-Vorpommern, DE

82, Institute of Clinical Chemistry and Laboratory Medicine, University Medicine Greifswald, Greifswald, Mecklenburg-Vorpommern, DE

83, Institute of Health and Biomedical Innovation, Queensland University of Technology, Brisbane, QLD, AU

84, Humus, Reykjavik, IS

85, Virginia Institute for Psychiatric & Behavioral Genetics, Virginia Commonwealth University, Richmond, VA, US

86, Clinical Genetics, Vrije Universiteit Medical Center, Amsterdam, NL

87, Complex Trait Genetics, Vrije Universiteit Amsterdam, Amsterdam, NL

88, Solid Biosciences, Boston, MA, US

89, Department of Psychiatry, Washington University in Saint Louis School of Medicine, Saint Louis, MO, US

90, Department of Biochemistry and Molecular Biology II, Institute of Neurosciences, Center for Biomedical Research, University of Granada, Granada, ES

91, Department of Psychiatry, University of Groningen, University Medical Center Groningen, Groningen, NL

92, Department of Psychiatry and Psychotherapy, Medical Center of the University of Munich, Campus Innenstadt, Munich, DE

93, Institute of Psychiatric Phenomics and Genomics (IPPG), Medical Center of the University of Munich, Campus Innenstadt, Munich, DE

94, Division of Cancer Epidemiology and Genetics, National Cancer Institute, Bethesda, MD, US

95, Behavioral Health Services, Kaiser Permanente Washington, Seattle, WA, US

96, Faculty of Medicine, Department of Psychiatry, University of Iceland, Reykjavik, IS

97, School of Medicine and Dentistry, James Cook University, Townsville, QLD, AU

98, Institute of Health and Wellbeing, University of Glasgow, Glasgow, GB

99, deCODE Genetics / Amgen, Reykjavik, IS

100, College of Biomedical and Life Sciences, Cardiff University, Cardiff, GB

101, Institute of Epidemiology and Social Medicine, University of Münster, Münster, Nordrhein-Westfalen, DE

102, Institute for Community Medicine, University Medicine Greifswald, Greifswald, Mecklenburg-Vorpommern, DE

103, Department of Psychiatry, University of California, San Diego, San Diego, CA, US

104, KG Jebsen Centre for Psychosis Research, Norway Division of Mental Health and Addiction, Oslo University Hospital, Oslo, NO

105, Medical Genetics Section, CGEM, IGMM, University of Edinburgh, Edinburgh, GB

106, Clinical Neurosciences, University of Cambridge, Cambridge, GB

107, Internal Medicine, Erasmus MC, Rotterdam, Zuid-Holland, NL

108, Roche Pharmaceutical Research and Early Development, Neuroscience, Ophthalmology and Rare Diseases Discovery & Translational Medicine Area, Roche Innovation Center Basel, F. Hoffmann-La Roche Ltd, Basel, CH

109, Department of Psychiatry and Psychotherapy, University Medicine Greifswald, Greifswald, Mecklenburg-Vorpommern, DE

110, Department of Psychiatry, Leiden University Medical Center, Leiden, NL

111, Virginia Institute for Psychiatric & Behavioral Genetics, Virginia Commonwealth University, Richmond, VA, US

112, Computational Sciences Center of Emphasis, Pfizer Global Research and Development, Cambridge, MA, US

113, Institute for Molecular Bioscience; Queensland Brain Institute, The University of Queensland, Brisbane, QLD, AU

114, Department of Psychiatry, University of Münster, Münster, Nordrhein-Westfalen, DE

115, Department of Psychiatry, Melbourne Medical School, University of Melbourne, Melbourne, AU

116, Institute of Medical Genetics and Pathology, University Hospital Basel, University of Basel, Basel, CH

117, Institute of Neuroscience and Medicine (INM-1), Research Center Juelich, Juelich, DE

118 Amsterdam Public Health Institute (APH), Vrije Universiteit Medical Center, Amsterdam, NL

119, Centre for Integrative Biology, Università degli Studi di Trento, Trento, Trentino-Alto Adige, IT

120, Department of Psychiatry and Psychotherapy, Medical Center, University of Freiburg, Faculty of Medicine, University of Freiburg, Freiburg, DE

121, Psychiatry, Kaiser Permanente Northern California, San Francisco, CA, US

122, Medical Research Council Human Genetics Unit, Institute of Genetics and Molecular Medicine, University of Edinburgh, Edinburgh, GB

123, Department of Psychiatry, University of Toronto, Toronto, ON, CA

124, Centre for Addiction and Mental Health, Toronto, ON, CA

125, Division of Psychiatry, University College London, London, GB

126, Neuroscience Therapeutic Area, Janssen Research and Development, LLC, Titusville, NJ, US

127, Institute of Molecular and Cell Biology, University of Tartu, Tartu, EE

128, Psychosis Research Unit, Aarhus University Hospital, Risskov, Aarhus, DK

129, University of Liverpool, Liverpool, GB

130, Mental Health Center Copenhagen, Copenhagen Universtity Hospital, Copenhagen, DK

131, Human Genetics and Computational Biomedicine, Pfizer Global Research and Development, Groton, CT, US

132, Psychiatry, Harvard Medical School, Boston, MA, US

133, Psychiatry, University of Iowa, Iowa City, IA, US

134, Department of Psychiatry and Behavioral Sciences, Johns Hopkins University, Baltimore, MD, US

135, Department of Psychiatry and Psychotherapy, University Medical Center Göttingen, Goettingen, Niedersachsen, DE

136, Human Genetics Branch, NIMH Division of Intramural Research Programs, Bethesda, MD, US

137, Faculty of Medicine, University of Iceland, Reykjavik, IS

138, Child and Adolescent Psychiatry, Erasmus MC, Rotterdam, Zuid-Holland, NL

139, Psychiatry, Erasmus MC, Rotterdam, Zuid-Holland, NL

140, Psychiatry, Dalhousie University, Halifax, NS, CA

141, Division of Epidemiology, New York State Psychiatric Institute, New York, NY, US

142, Department of Clinical Medicine, University of Copenhagen, Copenhagen, DK

143, Department of Medical & Molecular Genetics, King's College London, London, GB

144, Psychiatry & Behavioral Sciences, Stanford University, Stanford, CA, US

145, NIHR Maudsley Biomedical Research Centre, King's College London, London, GB

146, Genetics, University of North Carolina at Chapel Hill, Chapel Hill, NC, US

147, Psychiatry, University of North Carolina at Chapel Hill, Chapel Hill, NC, US

MEGASTROKE CONSORTIUM

Rainer Malik 1, Ganesh Chauhan 2, Matthew Traylor 3, Muralidharan Sargurupremraj 4,5, Yukinori Okada 6,7,8, Aniket Mishra 4,5, Loes Rutten-Jacobs 3, Anne-Katrin Giese 9, Sander W van der Laan 10, Solveig Gretarsdottir 11, Christopher D Anderson 12,13,14,14, Michael Chong 15, Hieab HH Adams 16,17, Tetsuro Ago 18, Peter Almgren 19, Philippe Amouyel 20,21, Hakan Ay 22,13, Traci M Bartz 23, Oscar R Benavente 24, Steve Bevan 25, Giorgio B Boncoraglio 26, Robert D Brown, Jr. 27, Adam S Butterworth 28,29, Caty Carrera 30,31, Cara L Carty 32,33, Daniel I Chasman 34,35, Wei-Min Chen 36, John W Cole 37, Adolfo Correa 38, Ioana Cotlarciuc 39, Carlos Cruchaga 40,41, John Danesh 28,42,43,44, Paul IW de Bakker 45,46, Anita L DeStefano 47,48, Marcel den Hoed 49, Qing Duan 50, Stefan T Engelter 51,52, Guido J Falcone 53,54, Rebecca F Gottesman 55, Raji P Grewal 56, Vilmundur Gudnason 57,58, Stefan Gustafsson 59, Jeffrey Haessler 60, Tamara B Harris 61, Ahamad Hassan 62, Aki S Havulinna 63,64, Susan R Heckbert 65, Elizabeth G Holliday 66,67, George Howard 68, Fang-Chi Hsu 69, Hyacinth I Hyacinth 70, M Arfan Ikram 16, Erik Ingelsson 71,72, Marguerite R Irvin 73, Xueqiu Jian 74, Jordi Jiménez-Conde 75, Julie A Johnson 76,77, J Wouter Jukema 78, Masahiro Kanai 6,7,79, Keith L Keene 80,81, Brett M Kissela 82, Dawn O Kleindorfer 82, Charles Kooperberg 60, Michiaki Kubo 83, Leslie A Lange 84, Carl D Langefeld 85, Claudia Langenberg 86, Lenore J Launer 87, Jin-Moo Lee 88, Robin Lemmens 89,90, Didier Leys 91, Cathryn M Lewis 92,93, Wei-Yu Lin 28,94, Arne G Lindgren 95,96, Erik Lorentzen 97, Patrik K Magnusson 98, Jane Maguire 99, Ani Manichaikul 36, Patrick F McArdle 100, James F Meschia 101, Braxton D Mitchell 100,102, Thomas H Mosley 103,104, Michael A Nalls 105,106, Toshiharu Ninomiya 107, Martin J O'Donnell 15,108, Bruce M Psaty 109,110,111,112, Sara L Pulit 113,45, Kristiina Rannikmäe 114,115, Alexander P Reiner 65,116, Kathryn M Rexrode 117, Kenneth Rice 118, Stephen S Rich 36, Paul M Ridker 34,35, Natalia S Rost 9,13, Peter M Rothwell 119, Jerome I Rotter 120,121, Tatjana Rundek 122, Ralph L Sacco 122, Saori Sakaue 7,123, Michele M Sale 124, Veikko Salomaa 63, Bishwa R Sapkota 125, Reinhold Schmidt 126, Carsten O Schmidt 127, Ulf Schminke 128, Pankaj Sharma 39, Agnieszka Slowik 129, Cathie LM Sudlow 114,115, Christian Tanislav 130, Turgut Tatlisumak 131,132, Kent D Taylor 120,121, Vincent NS Thijs 133,134, Gudmar Thorleifsson 11, Unnur Thorsteinsdottir 11, Steffen Tiedt 1, Stella Trompet 135, Christophe Tzourio 5,136,137, Cornelia M van Duijn 138,139, Matthew Walters 140, Nicholas J Wareham 86, Sylvia Wassertheil-Smoller 141, James G Wilson 142, Kerri L Wiggins 109, Qiong Yang 47, Salim Yusuf 15, Najaf Amin 16, Hugo S Aparicio 185,48, Donna K Arnett 186, John Attia 187, Alexa S Beiser 47,48, Claudine Berr 188, Julie E Buring 34,35, Mariana Bustamante 189, Valeria Caso 190, Yu-Ching Cheng 191, Seung Hoan Choi 192,48, Ayesha Chowhan 185,48, Natalia Cullell 31, Jean-François Dartigues 193,194, Hossein Delavaran 95,96, Pilar Delgado 195, Marcus Dörr 196,197, Gunnar Engström 19, Ian Ford 198, Wander S Gurpreet 199, Anders Hamsten 200,201, Laura Heitsch 202, Atsushi Hozawa 203, Laura Ibanez 204, Andreea Ilinca 95,96, Martin Ingelsson 205, Motoki Iwasaki 206, Rebecca D Jackson 207, Katarina Jood 208, Pekka Jousilahti 63, Sara Kaffashian 4,5, Lalit Kalra 209, Masahiro Kamouchi 210, Takanari Kitazono 211, Olafur Kjartansson 212, Manja Kloss 213, Peter J Koudstaal 214, Jerzy Krupinski 215, Daniel L Labovitz 216, Cathy C Laurie 118, Christopher R Levi 217, Linxin Li 218, Lars Lind 219, Cecilia M Lindgren 220,221, Vasileios Lioutas 222,48, Yong Mei Liu 223, Oscar L Lopez 224, Hirata Makoto 225, Nicolas Martinez-Majander 172, Koichi Matsuda 225, Naoko Minegishi 203, Joan Montaner 226, Andrew P Morris 227,228, Elena Muiño 31, Martina Müller-Nurasyid 229,230,231, Bo Norrving 95,96, Soichi Ogishima 203, Eugenio A Parati 232, Leema Reddy Peddareddygari 56, Nancy L Pedersen 98,233, Joanna Pera 129, Markus Perola 63,234, Alessandro Pezzini 235, Silvana Pileggi 236, Raquel Rabionet 237, Iolanda Riba-Llena 30, Marta Ribasés 238, Jose R Romero 185,48, Jaume Roquer 239,240, Anthony G Rudd 241,242, Antti-Pekka Sarin 243,244, Ralhan Sarju 199, Chloe Sarnowski 47,48, Makoto Sasaki 245, Claudia L Satizabal 185,48, Mamoru Satoh 245, Naveed Sattar 246, Norie Sawada 206, Gerli Sibolt 172, Ásgeir Sigurdsson 247, Albert Smith 248, Kenji Sobue 245, Carolina Soriano-Tárraga 240, Tara Stanne 249, O Colin Stine 250, David J Stott 251, Konstantin Strauch 229,252, Takako Takai 203, Hideo Tanaka 253,254, Kozo Tanno 245, Alexander Teumer 255, Liisa Tomppo 172, Nuria P Torres-Aguila 31, Emmanuel Touze 256,257, Shoichiro Tsugane 206, Andre G Uitterlinden 258, Einar M Valdimarsson 259, Sven J van der Lee 16, Henry Völzke 255, Kenji Wakai 253, David Weir 260, Stephen R Williams 261, Charles DA Wolfe 241,242, Quenna Wong 118, Huichun Xu 191, Taiki Yamaji 206, Dharambir K Sanghera 125,169,170, Olle Melander 19, Christina Jern 171, Daniel Strbian 172,173, Israel Fernandez-Cadenas 31,30, W T Longstreth, Jr 174,65, Arndt Rolfs 175, Jun Hata 107, Daniel Woo 82, Jonathan Rosand 12,13,14, Guillaume Pare 15, Jemma C Hopewell 176, Danish Saleheen 177, Kari Stefansson 11,178, Bradford B Worrall 179, Steven J Kittner 37, Sudha Seshadri 180,48, Myriam Fornage 74,181, Hugh S Markus 3, Joanna MM Howson 28, Yoichiro Kamatani 6,182, Stephanie Debette 4,5, Martin Dichgans 1,183,184

1 Institute for Stroke and Dementia Research (ISD), University Hospital, LMU Munich, Munich, Germany

2 Centre for Brain Research, Indian Institute of Science, Bangalore, India

3 Stroke Research Group, Division of Clinical Neurosciences, University of Cambridge, UK

4 INSERM U1219 Bordeaux Population Health Research Center, Bordeaux, France

5 University of Bordeaux, Bordeaux, France

6 Laboratory for Statistical Analysis, RIKEN Center for Integrative Medical Sciences, Yokohama, Japan

7 Department of Statistical Genetics, Osaka University Graduate School of Medicine, Osaka, Japan

8 Laboratory of Statistical Immunology, Immunology Frontier Research Center (WPI-IFReC), Osaka University, Suita, Japan.

9 Department of Neurology, Massachusetts General Hospital, Harvard Medical School, Boston, MA, USA

10 Laboratory of Experimental Cardiology, Division of Heart and Lungs, University Medical Center Utrecht, University of Utrecht, Utrecht,Netherlands

11 deCODE genetics/AMGEN inc, Reykjavik, Iceland

12 Center for Genomic Medicine, Massachusetts General Hospital (MGH), Boston, MA, USA

13 J. Philip Kistler Stroke Research Center, Department of Neurology, MGH, Boston, MA, USA

14 Program in Medical and Population Genetics, Broad Institute, Cambridge, MA, USA

15 Population Health Research Institute, McMaster University, Hamilton, Canada

16 Department of Epidemiology, Erasmus University Medical Center, Rotterdam, Netherlands

17 Department of Radiology and Nuclear Medicine, Erasmus University Medical Center, Rotterdam, Netherlands

18 Department of Medicine and Clinical Science, Graduate School of Medical Sciences, Kyushu University, Fukuoka, Japan

19 Department of Clinical Sciences, Lund University, Malmö, Sweden

20 Univ. Lille, Inserm, Institut Pasteur de Lille, LabEx DISTALZ-UMR1167, Risk factors and molecular determinants of aging-related diseases, F-59000 Lille, France

21 Centre Hosp. Univ Lille, Epidemiology and Public Health Department, F-59000 Lille, France

22 AA Martinos Center for Biomedical Imaging, Department of Radiology, Massachusetts General Hospital, Harvard Medical School, Boston, MA, USA

23 Cardiovascular Health Research Unit, Departments of Biostatistics and Medicine, University of Washington, Seattle, WA, USA

24 Division of Neurology, Faculty of Medicine, Brain Research Center, University of British Columbia, Vancouver, Canada

25 School of Life Science, University of Lincoln, Lincoln, UK

26 Department of Cerebrovascular Diseases, Fondazione IRCCS Istituto Neurologico "Carlo Besta", Milano, Italy

27 Department of Neurology, Mayo Clinic Rochester, Rochester, MN, USA

28 MRC/BHF Cardiovascular Epidemiology Unit, Department of Public Health and Primary Care, University of Cambridge, Cambridge, UK

29 The National Institute for Health Research Blood and Transplant Research Unit in Donor Health and Genomics, University of Cambridge, UK

30 Neurovascular Research Laboratory, Vall d'Hebron Institut of Research, Neurology and Medicine Departments-Universitat Autònoma de Barcelona, Vall d’Hebrón Hospital, Barcelona, Spain

31 Stroke Pharmacogenomics and Genetics, Fundacio Docència i Recerca MutuaTerrassa, Terrassa, Spain

32 Children's Research Institute, Children's National Medical Center, Washington, DC, USA

33 Center for Translational Science, George Washington University, Washington, DC, USA

34 Division of Preventive Medicine, Brigham and Women's Hospital, Boston, MA, USA

35 Harvard Medical School, Boston, MA, USA

36 Center for Public Health Genomics, Department of Public Health Sciences, University of Virginia, Charlottesville, VA, USA

37 Department of Neurology, University of Maryland School of Medicine and Baltimore VAMC, Baltimore, MD, USA

38 Departments of Medicine, Pediatrics and Population Health Science, University of Mississippi Medical Center, Jackson, MS, USA

39 Institute of Cardiovascular Research, Royal Holloway University of London, UK & Ashford and St Peters Hospital, Surrey UK

40 Department of Psychiatry,The Hope Center Program on Protein Aggregation and Neurodegeneration (HPAN),Washington University, School of Medicine, St. Louis, MO, USA

41 Department of Developmental Biology, Washington University School of Medicine, St. Louis, MO, USA

42 NIHR Blood and Transplant Research Unit in Donor Health and Genomics, Department of Public Health and Primary Care, University of Cambridge, Cambridge, UK

43 Wellcome Trust Sanger Institute, Wellcome Trust Genome Campus, Hinxton, Cambridge, UK

44 British Heart Foundation, Cambridge Centre of Excellence, Department of Medicine, University of Cambridge, Cambridge, UK

45 Department of Medical Genetics, University Medical Center Utrecht, Utrecht, Netherlands

46 Department of Epidemiology, Julius Center for Health Sciences and Primary Care, University Medical Center Utrecht, Utrecht, Netherlands

47 Boston University School of Public Health, Boston, MA, USA

48 Framingham Heart Study, Framingham, MA, USA

49 Department of Immunology, Genetics and Pathology and Science for Life Laboratory, Uppsala University, Uppsala, Sweden

50 Department of Genetics, University of North Carolina, Chapel Hill, NC, USA

51 Department of Neurology and Stroke Center, Basel University Hospital, Switzerland

52 Neurorehabilitation Unit, University and University Center for Medicine of Aging and Rehabilitation Basel, Felix Platter Hospital, Basel, Switzerland

53 Department of Neurology, Yale University School of Medicine, New Haven, CT, USA

54 Program in Medical and Population Genetics, The Broad Institute of Harvard and MIT, Cambridge, MA, USA

55 Department of Neurology, Johns Hopkins University School of Medicine, Baltimore, MD, USA

56 Neuroscience Institute, SF Medical Center, Trenton, NJ, USA

57 Icelandic Heart Association Research Institute, Kopavogur, Iceland

58 University of Iceland, Faculty of Medicine, Reykjavik, Iceland

59 Department of Medical Sciences, Molecular Epidemiology and Science for Life Laboratory, Uppsala University, Uppsala, Sweden

60 Division of Public Health Sciences, Fred Hutchinson Cancer Research Center, Seattle, WA, USA

61 Laboratory of Epidemiology and Population Science, National Institute on Aging, National Institutes of Health, Bethesda, MD, USA

62 Department of Neurology, Leeds General Infirmary, Leeds Teaching Hospitals NHS Trust, Leeds, UK

63 National Institute for Health and Welfare, Helsinki, Finland

64 FIMM - Institute for Molecular Medicine Finland, Helsinki, Finland

65 Department of Epidemiology, University of Washington, Seattle, WA, USA

66 Public Health Stream, Hunter Medical Research Institute, New Lambton, Australia

67 Faculty of Health and Medicine, University of Newcastle, Newcastle, Australia

68 School of Public Health, University of Alabama at Birmingham, Birmingham, AL, USA

69 Department of Biostatistical Sciences, Wake Forest School of Medicine, Winston-Salem, NC, USA

70 Aflac Cancer and Blood Disorder Center, Department of Pediatrics, Emory University School of Medicine, Atlanta, GA, USA

71 Department of Medicine, Division of Cardiovascular Medicine, Stanford University School of Medicine, CA, USA

72 Department of Medical Sciences, Molecular Epidemiology and Science for Life Laboratory, Uppsala University, Uppsala, Sweden

73 Epidemiology, School of Public Health, University of Alabama at Birmingham, USA

74 Brown Foundation Institute of Molecular Medicine, University of Texas Health Science Center at Houston, Houston, TX, USA

75 Neurovascular Research Group (NEUVAS), Neurology Department, Institut Hospital del Mar d'Investigació Mèdica, Universitat Autònoma de Barcelona, Barcelona, Spain

76 Department of Pharmacotherapy and Translational Research and Center for Pharmacogenomics, University of Florida, College of Pharmacy, Gainesville, FL, USA

77 Division of Cardiovascular Medicine, College of Medicine, University of Florida, Gainesville, FL, USA

78 Department of Cardiology, Leiden University Medical Center, Leiden, the Netherlands

79 Program in Bioinformatics and Integrative Genomics, Harvard Medical School, Boston, MA, USA

80 Department of Biology, East Carolina University, Greenville, NC, USA

81 Center for Health Disparities, East Carolina University, Greenville, NC, USA

82 University of Cincinnati College of Medicine, Cincinnati, OH, USA

83 RIKEN Center for Integrative Medical Sciences, Yokohama, Japan

84 Department of Medicine, University of Colorado Denver, Anschutz Medical Campus, Aurora, CO, USA

85 Center for Public Health Genomics and Department of Biostatistical Sciences, Wake Forest School of Medicine, Winston-Salem, NC, USA

86 MRC Epidemiology Unit, University of Cambridge School of Clinical Medicine, Institute of Metabolic Science, Cambridge Biomedical Campus, Cambridge, UK

87 Intramural Research Program, National Institute on Aging, National Institutes of Health, Bethesda, MD, USA

88 Department of Neurology, Radiology, and Biomedical Engineering, Washington University School of Medicine, St. Louis, MO, USA

89 KU Leuven – University of Leuven, Department of Neurosciences, Experimental Neurology, Leuven, Belgium

90 VIB Center for Brain & Disease Research, University Hospitals Leuven, Department of Neurology, Leuven, Belgium

91 Univ.-Lille, INSERM U 1171. CHU Lille. Lille, France

92 Department of Medical and Molecular Genetics, King's College London, London, UK

93 SGDP Centre, Institute of Psychiatry, Psychology & Neuroscience, King's College London, London, UK

94 Northern Institute for Cancer Research, Paul O'Gorman Building, Newcastle University, Newcastle, UK

95 Department of Clinical Sciences Lund, Neurology, Lund University, Lund, Sweden

96 Department of Neurology and Rehabilitation Medicine, Skåne University Hospital, Lund, Sweden

97 Bioinformatics Core Facility, University of Gothenburg, Gothenburg, Sweden

98 Department of Medical Epidemiology and Biostatistics, Karolinska Institutet, Stockholm, Sweden

99 University of Technology Sydney, Faculty of Health, Ultimo, Australia

100 Department of Medicine, University of Maryland School of Medicine, MD, USA

101 Department of Neurology, Mayo Clinic, Jacksonville, FL, USA

102 Geriatrics Research and Education Clinical Center, Baltimore Veterans Administration Medical Center, Baltimore, MD, USA

103 Division of Geriatrics, School of Medicine, University of Mississippi Medical Center, Jackson, MS, USA

104 Memory Impairment and Neurodegenerative Dementia Center, University of Mississippi Medical Center, Jackson, MS, USA

105 Laboratory of Neurogenetics, National Institute on Aging, National institutes of Health, Bethesda, MD, USA

106 Data Tecnica International, Glen Echo MD, USA

107 Department of Epidemiology and Public Health, Graduate School of Medical Sciences, Kyushu University, Fukuoka, Japan

108 Clinical Research Facility, Department of Medicine, NUI Galway, Galway, Ireland

109 Cardiovascular Health Research Unit, Department of Medicine, University of Washington, Seattle, WA, USA

110 Department of Epidemiology, University of Washington, Seattle, WA

111 Department of Health Services, University of Washington, Seattle, WA, USA

112 Kaiser Permanente Washington Health Research Institute, Seattle, WA, USA

113 Brain Center Rudolf Magnus, Department of Neurology, University Medical Center Utrecht, Utrecht, The Netherlands

114 Usher Institute of Population Health Sciences and Informatics, University of Edinburgh, Edinburgh, UK

115 Centre for Clinical Brain Sciences, University of Edinburgh, Edinburgh, UK

116 Fred Hutchinson Cancer Research Center, University of Washington, Seattle, WA, USA

117 Department of Medicine, Brigham and Women's Hospital, Boston, MA, USA

118 Department of Biostatistics, University of Washington, Seattle, WA, USA

119 Nuffield Department of Clinical Neurosciences, University of Oxford, UK

120 Institute for Translational Genomics and Population Sciences, Los Angeles Biomedical Research Institute at Harbor-UCLA Medical Center, Torrance, CA, USA

121 Division of Genomic Outcomes, Department of Pediatrics, Harbor-UCLA Medical Center, Torrance, CA, USA

122 Department of Neurology, Miller School of Medicine, University of Miami, Miami, FL, USA

123 Department of Allergy and Rheumatology, Graduate School of Medicine, the University of Tokyo, Tokyo, Japan

124 Center for Public Health Genomics, University of Virginia, Charlottesville, VA, USA

125 Department of Pediatrics, College of Medicine, University of Oklahoma Health Sciences Center, Oklahoma City, OK, USA

126 Department of Neurology, Medical University of Graz, Graz, Austria

127 University Medicine Greifswald, Institute for Community Medicine, SHIP-KEF, Greifswald, Germany

128 University Medicine Greifswald, Department of Neurology, Greifswald, Germany

129 Department of Neurology, Jagiellonian University, Krakow, Poland

130 Department of Neurology, Justus Liebig University, Giessen, Germany

131 Department of Clinical Neurosciences/Neurology, Institute of Neuroscience and Physiology, Sahlgrenska Academy at University of Gothenburg, Gothenburg, Sweden

132 Sahlgrenska University Hospital, Gothenburg, Sweden

133 Stroke Division, Florey Institute of Neuroscience and Mental Health, University of Melbourne, Heidelberg, Australia

134 Austin Health, Department of Neurology, Heidelberg, Australia

135 Department of Internal Medicine, Section Gerontology and Geriatrics, Leiden University Medical Center, Leiden, the Netherlands

136 INSERM U1219, Bordeaux, France

137 Department of Public Health, Bordeaux University Hospital, Bordeaux, France

138 Genetic Epidemiology Unit, Department of Epidemiology, Erasmus University Medical Center Rotterdam, Netherlands

139 Center for Medical Systems Biology, Leiden, Netherlands

140 School of Medicine, Dentistry and Nursing at the University of Glasgow, Glasgow, UK

141 Department of Epidemiology and Population Health, Albert Einstein College of Medicine, NY, USA

142 Department of Physiology and Biophysics, University of Mississippi Medical Center, Jackson, MS, USA

143 A full list of members and affiliations appears in the Supplementary Note

144 Department of Human Genetics, McGill University, Montreal, Canada

145 Department of Pathophysiology, Institute of Biomedicine and Translation Medicine, University of Tartu, Tartu, Estonia

146 Department of Cardiac Surgery, Tartu University Hospital, Tartu, Estonia

147 Clinical Gene Networks AB,Stockholm, Sweden

148 Department of Genetics and Genomic Sciences, The Icahn Institute for Genomics and Multiscale Biology Icahn School of Medicine at Mount Sinai, New York, NY , USA

149 Department of Pathophysiology, Institute of Biomedicine and Translation Medicine, University of Tartu, Biomeedikum, Tartu, Estonia

150 Integrated Cardio Metabolic Centre, Department of Medicine, Karolinska Institutet, Karolinska Universitetssjukhuset, Huddinge, Sweden.

151 Clinical Gene Networks AB, Stockholm, Sweden

152 Sorbonne Universités, UPMC Univ. Paris 06, INSERM, UMR_S 1166, Team Genomics & Pathophysiology of Cardiovascular Diseases, Paris, France

153 ICAN Institute for Cardiometabolism and Nutrition, Paris, France

154 Department of Biomedical Engineering, University of Virginia, Charlottesville, VA, USA

155 Group Health Research Institute, Group Health Cooperative, Seattle, WA, USA

156 Seattle Epidemiologic Research and Information Center, VA Office of Research and Development, Seattle, WA, USA

157 Cardiovascular Research Center, Massachusetts General Hospital, Boston, MA, USA

158 Department of Medical Research, Bærum Hospital, Vestre Viken Hospital Trust, Gjettum, Norway

159 Saw Swee Hock School of Public Health, National University of Singapore and National University Health System, Singapore

160 National Heart and Lung Institute, Imperial College London, London, UK

161 Department of Gene Diagnostics and Therapeutics, Research Institute, National Center for Global Health and Medicine, Tokyo, Japan

162 Department of Epidemiology, Tulane University School of Public Health and Tropical Medicine, New Orleans, LA, USA

163 Department of Cardiology,University Medical Center Groningen, University of Groningen, Netherlands

164 MRC-PHE Centre for Environment and Health, School of Public Health, Department of Epidemiology and Biostatistics, Imperial College London, London, UK

165 Department of Epidemiology and Biostatistics, Imperial College London, London, UK

166 Department of Cardiology, Ealing Hospital NHS Trust, Southall, UK

167 National Heart, Lung and Blood Research Institute, Division of Intramural Research, Population Sciences Branch, Framingham, MA, USA

168 A full list of members and affiliations appears at the end of the manuscript

169 Department of Phamaceutical Sciences, Collge of Pharmacy, University of Oklahoma Health Sciences Center, Oklahoma City, OK, USA

170 Oklahoma Center for Neuroscience, Oklahoma City, OK, USA

171 Department of Pathology and Genetics, Institute of Biomedicine, The Sahlgrenska Academy at University of Gothenburg, Gothenburg, Sweden

172 Department of Neurology, Helsinki University Hospital, Helsinki, Finland

173 Clinical Neurosciences, Neurology, University of Helsinki, Helsinki, Finland

174 Department of Neurology, University of Washington, Seattle, WA, USA

175 Albrecht Kossel Institute, University Clinic of Rostock, Rostock, Germany

176 Clinical Trial Service Unit and Epidemiological Studies Unit, Nuffield Department of Population Health, University of Oxford, Oxford, UK

177 Department of Genetics, Perelman School of Medicine, University of Pennsylvania, PA, USA

178 Faculty of Medicine, University of Iceland, Reykjavik, Iceland

179 Departments of Neurology and Public Health Sciences, University of Virginia School of Medicine, Charlottesville, VA, USA

180 Department of Neurology, Boston University School of Medicine, Boston, MA, USA

181 Human Genetics Center, University of Texas Health Science Center at Houston, Houston, TX, USA

182 Center for Genomic Medicine, Kyoto University Graduate School of Medicine, Kyoto, Japan

183 Munich Cluster for Systems Neurology (SyNergy), Munich, Germany

184 German Center for Neurodegenerative Diseases (DZNE), Munich, Germany

185 Boston University School of Medicine, Boston, MA, USA

186 University of Kentucky College of Public Health, Lexington, KY, USA

187 University of Newcastle and Hunter Medical Research Institute, New Lambton, Australia

188 Univ. Montpellier, Inserm, U1061, Montpellier, France

189 Centre for Research in Environmental Epidemiology, Barcelona, Spain

190 Department of Neurology, Università degli Studi di Perugia, Umbria, Italy

191 Department of Medicine, University of Maryland School of Medicine, Baltimore, MD, USA

192 Broad Institute, Cambridge, MA, USA

193 Univ. Bordeaux, Inserm, Bordeaux Population Health Research Center, UMR 1219, Bordeaux, France

194 Bordeaux University Hospital, Department of Neurology, Memory Clinic, Bordeaux, France

195 Neurovascular Research Laboratory. Vall d'Hebron Institut of Research, Neurology and Medicine Departments-Universitat Autònoma de Barcelona. Vall d’Hebrón Hospital, Barcelona, Spain

196 University Medicine Greifswald, Department of Internal Medicine B, Greifswald, Germany

197 DZHK, Greifswald, Germany

198 Robertson Center for Biostatistics, University of Glasgow, Glasgow, UK

199 Hero DMC Heart Institute, Dayanand Medical College & Hospital, Ludhiana, India

200 Atherosclerosis Research Unit, Department of Medicine Solna, Karolinska Institutet, Stockholm, Sweden

201 Karolinska Institutet, Stockholm, Sweden

202 Division of Emergency Medicine, and Department of Neurology, Washington University School of Medicine, St. Louis, MO, USA

203 Tohoku Medical Megabank Organization, Sendai, Japan

204 Department of Psychiatry, Washington University School of Medicine, St. Louis, MO, USA

205 Department of Public Health and Caring Sciences / Geriatrics, Uppsala University, Uppsala, Sweden

206 Epidemiology and Prevention Group, Center for Public Health Sciences, National Cancer Center, Tokyo, Japan

207 Department of Internal Medicine and the Center for Clinical and Translational Science, The Ohio State University, Columbus, OH, USA

208 Institute of Neuroscience and Physiology, the Sahlgrenska Academy at University of Gothenburg, Goteborg, Sweden

209 Department of Basic and Clinical Neurosciences, King's College London, London, UK

210 Department of Health Care Administration and Management, Graduate School of Medical Sciences, Kyushu University, Japan

211 Department of Medicine and Clinical Science, Graduate School of Medical Sciences, Kyushu University, Japan

212 Landspitali National University Hospital, Departments of Neurology & Radiology, Reykjavik, Iceland

213 Department of Neurology, Heidelberg University Hospital, Germany

214 Department of Neurology, Erasmus University Medical Center

215 Hospital Universitari Mutua Terrassa, Terrassa (Barcelona), Spain

216 Albert Einstein College of Medicine, Montefiore Medical Center, New York, NY, USA

217 John Hunter Hospital, Hunter Medical Research Institute and University of Newcastle, Newcastle, NSW, Australia

218 Centre for Prevention of Stroke and Dementia, Nuffield Department of Clinical Neurosciences, University of Oxford, UK

219 Department of Medical Sciences, Uppsala University, Uppsala, Sweden

220 Genetic and Genomic Epidemiology Unit, Wellcome Trust Centre for Human Genetics, University of Oxford, Oxford, UK

221 The Wellcome Trust Centre for Human Genetics, Oxford, UK

222 Beth Israel Deaconess Medical Center, Boston, MA, USA

223 Wake Forest School of Medicine, Wake Forest, NC, USA

224 Department of Neurology, University of Pittsburgh, Pittsburgh, PA, USA

225 BioBank Japan, Laboratory of Clinical Sequencing, Department of Computational biology and medical Sciences, Graduate school of Frontier Sciences, The University of Tokyo, Tokyo, Japan

226 Neurovascular Research Laboratory, Vall d'Hebron Institut of Research, Neurology and Medicine Departments-Universitat Autònoma de Barcelona. Vall d’Hebrón Hospital, Barcelona, Spain

227 Department of Biostatistics, University of Liverpool, Liverpool, UK

228 Wellcome Trust Centre for Human Genetics, University of Oxford, Oxford, UK

229 Institute of Genetic Epidemiology, Helmholtz Zentrum München - German Research Center for Environmental Health, Neuherberg, Germany

230 Department of Medicine I, Ludwig-Maximilians-Universität, Munich, Germany

231 DZHK (German Centre for Cardiovascular Research), partner site Munich Heart Alliance, Munich, Germany

232 Department of Cerebrovascular Diseases, Fondazione IRCCS Istituto Neurologico “Carlo Besta”, Milano, Italy

233 Karolinska Institutet, MEB, Stockholm, Sweden

234 University of Tartu, Estonian Genome Center, Tartu, Estonia, Tartu, Estonia

235 Department of Clinical and Experimental Sciences, Neurology Clinic, University of Brescia, Italy

236 Translational Genomics Unit, Department of Oncology, IRCCS Istituto di Ricerche Farmacologiche Mario Negri, Milano, Italy

237 Department of Genetics, Microbiology and Statistics, University of Barcelona, Barcelona, Spain

238 Psychiatric Genetics Unit, Group of Psychiatry, Mental Health and Addictions, Vall d’Hebron Research Institute (VHIR), Universitat Autònoma de Barcelona, Biomedical Network Research Centre on Mental Health (CIBERSAM), Barcelona, Spain

239 Department of Neurology, IMIM-Hospital del Mar, and Universitat Autònoma de Barcelona, Spain

240 IMIM (Hospital del Mar Medical Research Institute), Barcelona, Spain

241 National Institute for Health Research Comprehensive Biomedical Research Centre, Guy's & St. Thomas' NHS Foundation Trust and King's College London, London, UK

242 Division of Health and Social Care Research, King's College London, London, UK

243 FIMM-Institute for Molecular Medicine Finland, Helsinki, Finland

244 THL-National Institute for Health and Welfare, Helsinki, Finland

245 Iwate Tohoku Medical Megabank Organization, Iwate Medical University, Iwate, Japan

246 BHF Glasgow Cardiovascular Research Centre, Faculty of Medicine, Glasgow, UK

247 deCODE Genetics/Amgen, Inc., Reykjavik, Iceland

248 Icelandic Heart Association, Reykjavik, Iceland

249 Institute of Biomedicine, the Sahlgrenska Academy at University of Gothenburg, Goteborg, Sweden

250 Department of Epidemiology, University of Maryland School of Medicine, Baltimore, MD, USA

251 Institute of Cardiovascular and Medical Sciences, Faculty of Medicine, University of Glasgow, Glasgow, UK

252 Chair of Genetic Epidemiology, IBE, Faculty of Medicine, LMU Munich, Germany

253 Division of Epidemiology and Prevention, Aichi Cancer Center Research Institute, Nagoya, Japan

254 Department of Epidemiology, Nagoya University Graduate School of Medicine, Nagoya, Japan

255 University Medicine Greifswald, Institute for Community Medicine, SHIP-KEF, Greifswald, Germany

256 Department of Neurology, Caen University Hospital, Caen, France

257 University of Caen Normandy, Caen, France

258 Department of Internal Medicine, Erasmus University Medical Center, Rotterdam, Netherlands

259 Landspitali University Hospital, Reykjavik, Iceland

260 Survey Research Center, University of Michigan, Ann Arbor, MI, USA

261 University of Virginia Department of Neurology, Charlottesville, VA, USA

1.	Hall LS, Adams MJ, Arnau-Soler A, et al. Genome-wide meta-analyses of stratified depression in Generation Scotland and UK Biobank. *Translational Psychiatry.* 2018;8(1):9.
